# NetREm Network Regression Embeddings reveal cell-type transcription factor coordination for gene regulation

**DOI:** 10.1101/2023.10.25.563769

**Authors:** Saniya Khullar, Xiang Huang, Raghu Ramesh, John Svaren, Daifeng Wang

## Abstract

Transcription factor (TF) coordination plays a key role in target gene (TG) regulation via protein-protein interactions (PPIs) and DNA co-binding to regulatory elements. Single-cell technologies facilitate gene expression measurement for individual cells and cell-type identification, yet the connection between TF coordination and TG regulation of various cell types remains unclear. To address this, we have developed a novel computational approach, Network Regression Embeddings (NetREm), to reveal cell-type TF-TF coordination activities for TG regulation. NetREm leverages network-constrained regularization using prior knowledge of direct and/or indirect PPIs among TFs to analyze single-cell gene expression data. We test NetREm by simulation data and benchmark its performance in 4 real-world applications that have gold standard TF-TG networks available: mouse (mESCs) and simulated human (hESCs) embryonic stem (ESCs), human hematopoietic stem (HSCs), and mouse dendritic (mDCs) cells. Further, we showcase NetREm’s ability to prioritize valid novel TF-TF coordination links in human Peripheral Blood Mononuclear cell (PBMC) sub-types. We apply NetREm to analyze various cell types in both central (CNS) and peripheral (PNS) nerve system (NS) (e.g. neuronal, glial, Schwann cells (SCs)) as well as in Alzheimer’s disease (AD). Our findings uncover cell-type coordinating TFs and identify new TF-TG candidate links. We validate our top predictions using Cut&Run and knockout loss-of-function expression data in rat/mouse models and compare results with additional functional genomic data, including expression quantitative trait loci (eQTL) and Genome-Wide Association Studies (GWAS) to link genetic variants (single nucleotide polymorphisms (SNPs)) to TF coordination. NetREm is open-source available on GitHub as a software package.

## Introduction

Transcription factors (TFs) collaborate to activate or repress the expression of their target genes (TGs) by binding to specific DNA sequences known as TF binding sites (TFBSs) on regulatory elements (REs; enhancers, promoters). These REs are shaped by the combinatorial interactions of multiple TFs, forming transcriptional regulatory modules (TRMs) that govern transcription initiation (Guo and Gifford 2017; Wunderlich and Mirny 2009). TFs often work in concert, either as stable complexes or by enhancing the binding affinity of nearby TFs (Nie et al. 2020; Zhao 2023). Despite the common occurrence of TF-TF coordination, the complexities underlying this phenomenon are not yet fully understood (Ibarra et. al 2020). The coordinated activity of core TFs, among 100s expressed, defines a cell type’s unique transcriptional landscape and identity (Lee et al. 2012; D’Alessio et al. 2015). Indeed, the TF binding grammar is complex and context-specific, influenced by proteins occupying shared REs and the chromatin structure (Bentsen et. al 2022).

Gene regulatory networks (GRNs: models of TG regulation by TFs) must incorporate the complex combinatorial coordination (nature: cooperativity, antagonism; type: direct, indirect) among TFs, which is facilitated by the positioning of TFBSs within REs (Berenson et al. 2023; Ouyang et al. 2009). Historically, classical models of TF cooperativity were predicated on direct protein-protein interactions (PPIs) since many TFs may form complexes (to increase their binding affinity and motif specificity (Sönmezer et al. 2021)) to co-regulate TGs (Wang et al. 2023). Based on recent studies, it is now understood that TFs may jointly coordinate TG transcription even without direct PPIs (Rao et al. 2021; Wang et al. 2009). Pioneer TFs, for instance, recognize and bind to their TFBSs even within closed chromatin, inducing local chromatin remodeling (e.g., nucleosome eviction), thereby exposing TFBSs and rendering DNA more amenable to subsequent binding by settler TFs (Sinha et al. 2023; Mayran and Drouin 2018). Tethered-binding mechanisms, involving coactivators/corepressors, allow TFs to regulate TGs cooperatively by recruiting intermediary proteins (e.g., p300/CBP, mediator complexes) (Spitz and Furlong 2012; Ibarra et. al 2020). Antagonistic coordination, via sequestration and/or competition for TFBSs, generates cell-state heterogeneity by driving opposing effects on epigenetic programs and TG expression; some TFs bind DNA only in the presence of cooperating TFs and absence of antagonistic TFs (Hu et al. 2022; Berenson et al. 2023).

Despite extensive single-cell data analyses (Mathys et. al 2019; Wang et. al 2018), the extent of TF-TF coordination in regulating TGs across various cell types remains unclear (Nie et al. 2020; Ibarra et. al 2020; Karczewski et al. 2011; Hannenhalli and Levy 2002). Understanding this is crucial for dissecting GRNs driving cognitive, motor, and behavioral processes, often dysregulated in disease states. GRN-inference tools often reverse-engineer expression data to find coordinated patterns and gene-gene interactions, uncovering meaningful biological signals from noise (Wang et al. 2023; Campos et al. 2019). State-of-the-art (SOTA) tools (e.g. BEELINE(Pratapa et. al 2020), SCENIC(Aibar et. al 2017), GRNBoost(Moerman et al. 2019), Signac(Stuart et. al 2021), scGRNom(Jin et al. 2021)) create cell-type GRNs using various methods (e.g. co-expression, correlation *r*, info theory, machine learning, multi-omics integration) but often overlook interdependent TF-TF PPI networks (PPINs) crucial for TG regulation.

Neglecting PPINs may lead these tools to disregard *TF*_*i*_-TG pairs with uncorrelated expression (Zaborowski and Walther 2020), which may bury true biological relations, as *TF*_*i*_ could still regulate the TG through complex avenues, often coordinating synergistically or antagonistically with other TFs. Coordinating TFs are often co-expressed (Ouyang et al. 2009) and amplify noise in expression profiles (Nagamine et al. 2005; Perna et al. 2020; Parab et al. 2022). Without additional context, GRN-inference tools might remove truly causal correlated TFs that co-regulate TGs, excluding fundamental aspects of GRN mechanisms (Nicodemus and Malley 2009). Thus, these tools ought to incorporate various levels of expression regulation like PPIs for TFs as they can help reveal these dynamics of TF-TF coordination (Nagamine et al. 2005; Perna et al. 2020; Parab et al. 2022). Tools like BGRMI (Iglesias-Martinez et al. 2016) identify relevant TF complexes using PPIs but ignore inherent PPIN structures and indirect PPIs. RTNduals (Chagas et al. 2019) predicts TF co-regulatory behavior solely from expression data, which may not always yield outputs. SCINET (Mohammadi et. al 2019) reconstructs cell-type interactomes, while TF-Cluster (Nie et al. 2011) identifies coordinated TFs but avoids using prior info. Neither tool focuses on TG regulation.

Previous studies leveraged network-regularized (e.g. graph-regularized, network Lasso) regression models to identify disease-associated genes and networks, incorporating biological info as metadata (Li and Li 2008; Wang et al. 2015; Dirmeier et al. 2018; Li and Li 2010; Kim et al. 2013). These models, guided and constrained by prior knowledge, enhance their biological relevance; however, they are not tailored to elucidate TF-TF coordination in TG regulation. There is a growing need for these models to advance, assimilating info from regression data and prior network inputs to generate robust embeddings that may extract significant relationships, offering new insights in biology and beyond (Chu et al. 2023; Gharavi et al. 2021; Choy et al. 2019).

To bridge these gaps, we developed NetREm (Network Regression Embeddings), a novel computational framework to infer cell-type TF-TF coordination activities for TG regulation. Building on established network-regularized regression techniques, NetREm integrates multimodal data (e.g. TF binding profiles, direct/indirect TF-TF PPIs, derived TF-TF colocalization, gene expression, chromatin interaction, scATAC-seq epigenomic markers) capturing intricate aspects of TG regulation. It constructs robust predictive models for TF-TG regulation (complementary GRNs) as well as TF-TF coordination. A distinct feature of NetREm is its innovative ability to generate network regression embeddings, which identify and quantify coordination among cell-type TFs for co-regulating individual TGs. Public databases like STRINGdb provide direct and/or indirect, organism-specific, global, undirected, cell-type-agnostic PPIs for >12k organisms (Szklarczyk et al. 2023). Despite PPINs having some incorrect PPIs (False Positives (FPs)) and being largely incomplete (Kotlyar et. al 2022), it is helpful to integrate TF-TF PPINs as prior info (McCalla et al. 2023; Li and Jackson 2015; Ghanbari et al. 2015; Imoto et al. 2003; Mukherjee and Speed 2008). This improves NetREm’s predicted GRNs from expression data. NetREm also addresses the need for not only cell-type-specific annotation of known PPIs, but also discovery of new PPIs. It provides much-needed insights into TRMs, reveals cell-type- and disease-specific TF-TF PPINs, and aids interactome studies in uncovering disease gene properties and differential PPIN rewiring (Göös et al. 2022; Sevimoglu and Arga 2014). To demonstrate, we apply NetREm to simulation data and NS cell types: myelinating (mSCs) vs. non-myelinating (nmSCs) SCs, as well as in Alzheimer’s disease (AD) vs. control states in neurons/glia. NetREm is an open-source tool for general-purpose use.

## Results

### Overview of Network Regression Embeddings (NetREm)

We developed NetREm as a robust, multi-omics, computational approach to build and integrate networks of TF-to-TG regulation and TF-TF coordination in a cell-type-specific manner (**Fig. 1**). NetREm is for applications where predictors form subnetworks to influence the outcome (Li and Liu 2022). NetREm uses single-cell or bulk gene expression data, a comprehensive PPIN (e.g. STRINGdb) that globally captures verified direct/indirect functional associations among TFs (Peng et al. 2017). Providing a prior, candidate GRN (e.g. initial TF-RE-TG links) is optional but recommended as it improves biological relevance of outputs with its initial feature selection. NetREm integrates this multi-modal data, applying a 2-step optimization process for each TG: **1**. It formulates a network-regularized regression problem, using the input PPIN as a priori info, to sift through *N* candidate TFs, to find those most likely to co-regulate TG. **2**. To solve this regression problem, it then employs Singular Value Decomposition (SVD) to create network regression embeddings, which are used in Lasso regression to predict TG expression. In the process, NetREm learns not only key TFs (coefficients *c*^*^, a complementary GRN) but also a TG-specific TF-TF coordination network - 100 ≤ *B* ≤ 100, predicting cooperative (> 0) or antagonistic (< 0) TF-TF relations for regulating TG. It also annotates known PPIs, uncovering confident cell-type-specific TF-TF PPI subnetworks involved in TG regulation (Szklarczyk et al. 2023). By integrating genomic and foundational PPIs, NetREm uses transfer learning to discover unanticipated, biologically significant TF-TG links and novel functional TF-TF relations for future investigation (Zhang et al. 2013). Aggregating outputs across all TGs, yields cell-type-specific directed TF-TG regulatory and undirected TF-TF coordination (-100 ≤ 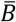 ≤100) networks. These outputs support downstream analyses, like linking non-coding SNPs to potential regulatory roles via expression trait loci (eQTL) SNP-TG (i.e. eSNP-eTG) associations in our networks.

**Figure 1.**
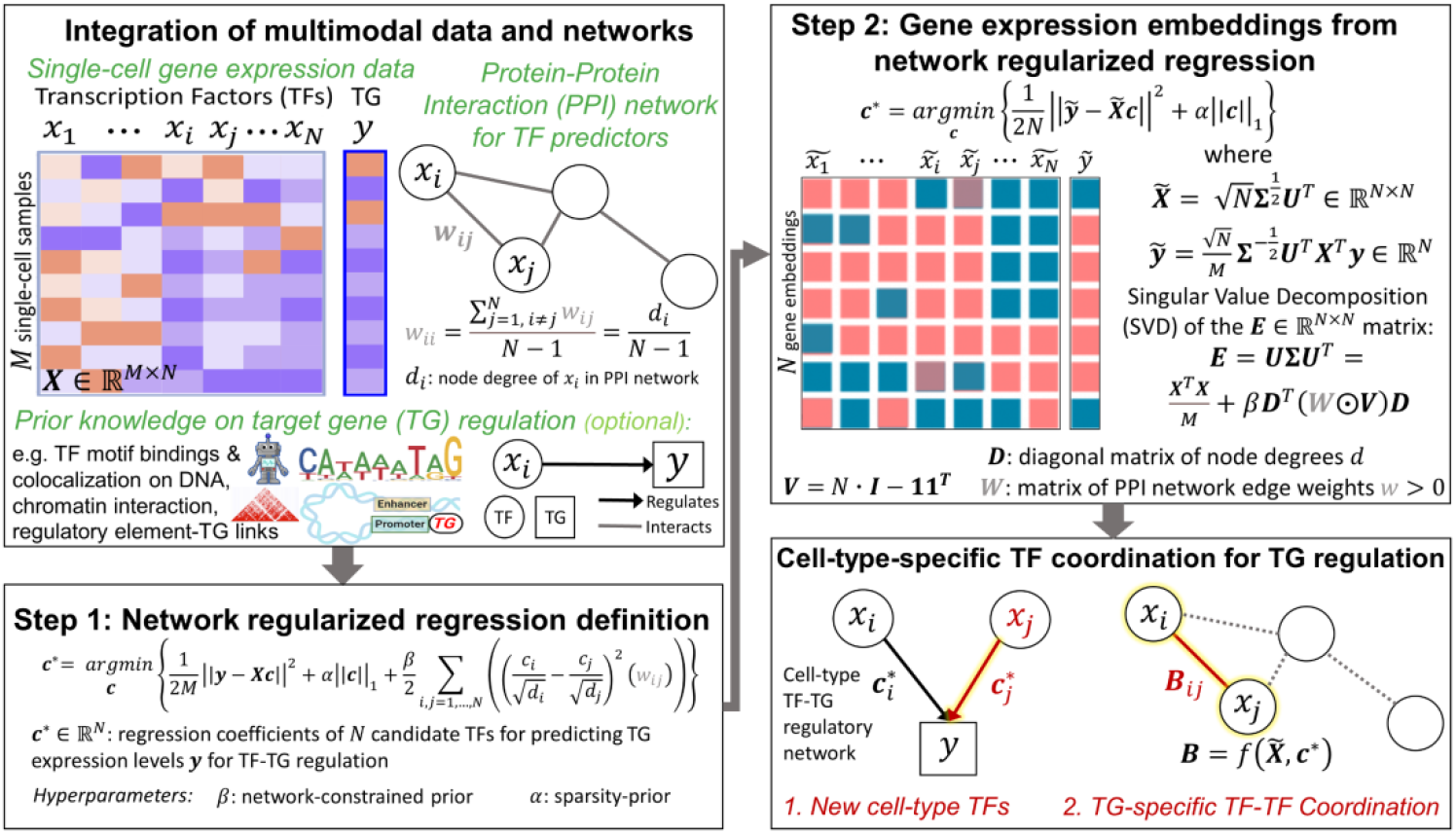
Overview of Network Regression Embeddings (NetREm) a multi-step, multi-omics computational framework to construct comprehensive cell-type-specific TF-TG regulatory and TF-TF coordination 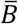 networks. We apply this pipeline for each TG in a cell-type. NetREm integrates multimodal data: gene expression (*X* ∈ ℝ^*M* × *N*^, *y* ∈ ℝ^*M*^) for *M* cells (samples), direct/indirect TF-TF PPIs with weights *W*, optional prior GRN information (e.g. TF-RE-TG links). This prior GRN helps select only relevant TF predictors for TG (from multiomics data) to reduce dimensionality of *X*: *M* (rows), *N* TFs (columns). *X* and *y* are standardized (mean *μ*: 0, standard deviation *σ*:1) across cells for TFs and TG. Goal: identify TFs, out of *N* candidates, whose expressions best predict TG expression *y*. **Step 1** sets up a PPIN-regularized regression problem to identify optimal TFs for a TG, guided by network *β*, sparsity *α* hyperparameters. **Step 2** solves this, transforming (*X, y*) and PPIs to latent space gene expression embeddings 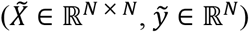 by SVD on an 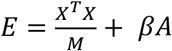 matrix (*A* = *D*^*T*^(*W⨀V*)*D*). *E* combines expression relations and PPIN information. 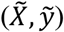 undergo Lasso regression (via model-type: Lasso (*α* is given) or LassoCV (*α* chosen by cross-validation (CV); default: no intercept) to predict optimal coefficients *c*^*^ ∈ ℝ^*N*^ for TG. NetREm outputs 2 networks capturing different aspects of TG regulation that can be integrated. #1: links optimal TFs to TG by *c*^*^ (> 0: activator, < 0: repressor), in a directed TF-TG regulatory network, a complementary GRN. This likely uncovers novel cell-type TFs like *x*_*j*_ and reflects underlying biology of coordination among TFs. If we input a prior GRN, we may use it to help annotate our TF-TG links with biological metadata (e.g. TF-RE-TG network). <u>#2</u>: TG-specific undirected TF-TF coordination network *B* predicts indirect/direct relations among TFs to regulate TG. *TF*_*i*_ − *TF*_*j*_ coordination *B*_*ij*_ shows cooperative (> 0 if both are co-activators or co-repressors) or antagonistic: (< 0 if 1 is repressor, other is activator) co-regulation of TG. *B* is a matrix of *c*^*^-aware cosine similarity scores (**Methods**), a function of 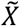 and *c*^*^. NetREm thus identifies novel coordination among cell-type TFs for co-regulating TG. Results are stitched together across runs for all TGs to obtain final cell-type outputs.

We demonstrate NetREm’s versatility across diverse real-world scenarios by benchmarking our TF-TG networks against established gold standard GRNs in human (hESCs, HSCs) and mouse (mESCs, mDCs) models. We use diverse techniques to assess 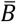 across these settings, as well as additional human contexts: 9 PBMCs, pooled stem cells from GTEx (Eraslan et. al 2022), 4 CNS cell types (Lake et al. 2018). Further, we highlight NetREm’s use of prior input GRNs, derived from multi-omics data, for specific contexts (cell types, diseases) in 2 human applications: 1. mSCs vs. nmSCs; 2. AD vs. controls in 8 neuron/glia cell types. The resulting TF-TG networks, integrated with TF-RE-TG annotations (**Methods**), yield enhanced, context-specific TF-RE-TG regulatory and coordination 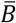 networks.

### Simulation study

We test NetREm on simulated single-cell gene expression data for 10,000 cells as a proof-of-concept, where: *X* (matrix of expression levels for *N* = 5 candidate TF predictors) and *y* (TG expression vector) are drawn from a normal distribution with dropouts to achieve ∼40% sparsity, mimicking single-cell data (**Fig. S1A**). Train data (70%; *M*: 7,000 cells) standardizes (*X*_*train*_, *y*_*train*_, *X*_*test*_, *y*_*test*_) so each variable has (*μ* = 0, *σ* = 1). TFs 1 to 5 have expression with Pearson *r*(*TF, TG*) ≈ [0.9, 0.5, 0.4, -0.3, -0.8] with *y*, respectively, in train and test data (**Fig. 2A, Fig. S1B-C**). Our PPIN of known direct and/or indirect PPIs among TFs has strong *w* for *TF*_1_ − *TF*_2_ (0.8), *TF*_4_ − *TF*_5_ (0.95). We set *w* to η = 0.01 for missing PPIs, making our PPIN fully-connected. NetREm(*β* = 1, *α* = 0.1) predicts *y* based on *X*, subject to the PPIN constraint. It outputs 2 networks: TF-TG regulation (*c*^*^) relates expression levels of TFs to TG expression (**Fig. 2B**) and TG-specific TF-TF coordination (*B* scores) (**Fig. 2C**) of behavior among TFs to regulate TG.

**Figure 2.**
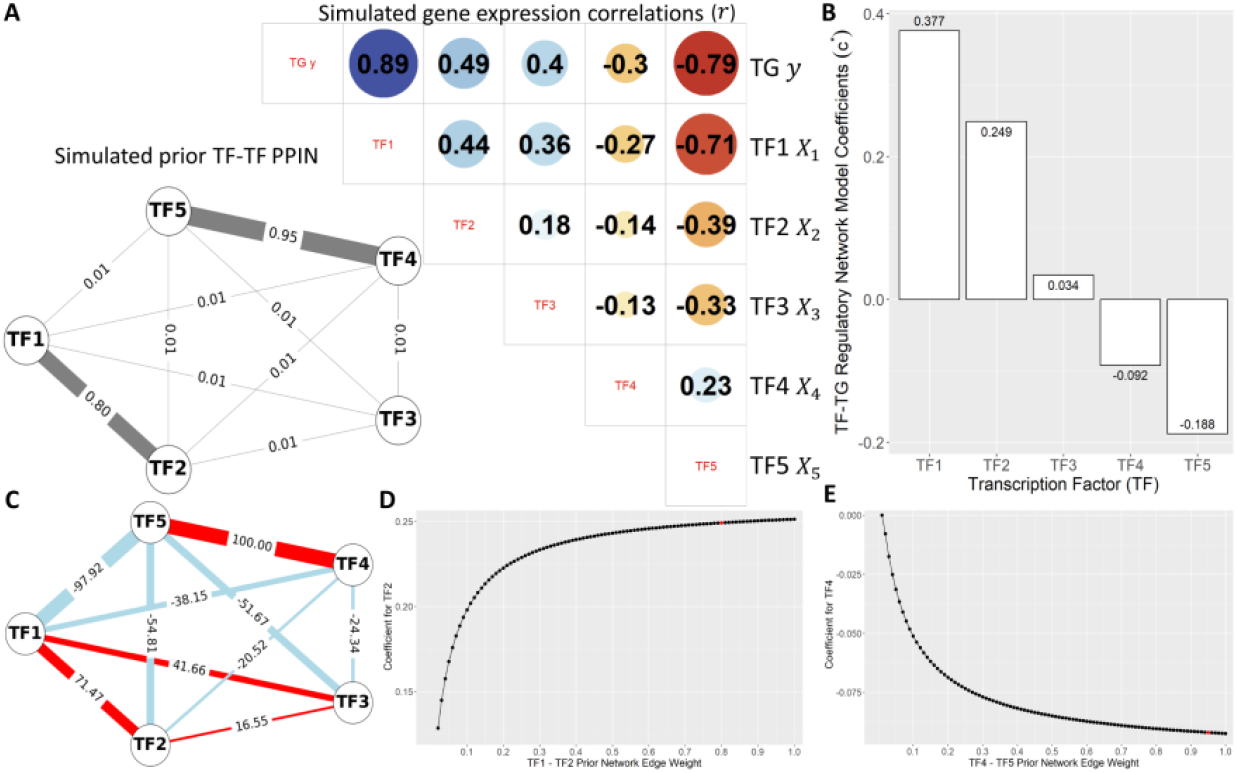
Simulation study of 5 TFs, 1 TG **A)** Bottom left: prior, undirected TF-TF PPIN with default weights (η = 0.01); strong experimentally-verified connections: *TF*_1_-*TF*_2_ (0.8); *TF*_4_-*TF*_5_ (0.95). Top right: Pearson *r* matrix in training data: *r*(*TF, TG*) ≈ [0.9, 0.5, 0.4, -0.3, -0.8]. Dot sizes: |*r*|: + (blue) or - (red) *r*. **B)** Coefficients *c*^*^ for TFs in TF-TG regulatory network, based on NetREm(*β* =1, *α* = 0.1, no intercept). Potential activators: *TF*_1_ to *TF*_3_. Repressors: *TF*_4_ and *TF*_5_. **C)** TG-specific TF-TF coordination network (scores: *B*) with red (antagonistic: -; activator-repressor links) and blue (cooperative: +; links between co-activators or co-repressors). Functionally-valid coordination: *TF*_1_-*TF*_2_ and *TF*_4_-*TF*_5_; others are novel. **D)** Effects of varying *TF*_1_-*TF*_2_ *w* in original PPIN, from 0.01 to 1 in 0.01 increments, holding all else fixed. Respective *TF*_2_ *c*^*^ increases monotonically in arc shape from 0.106 to 0.251. Red dot: PPIN *w*. **E)** Similar sensitivity analysis shows *TF*_4_’s |*c*^*^| increases, becoming more negative from 0 to -9.2e-2, as *TF*_4_-*TF*_5_ edge weight is perturbed.

We compare NetREm to 4 default Scikit-Learn (Pedregosa et al. 2011) benchmark regression models (BRMs) fit with no intercept: Linear Regression and 3 regularization ones (ElasticNetCV, LassoCV, RidgeCV). TFs with lower ranks have higher |*c*^*^| and are more important. Absolute values of *c*^*^ for *TF*_2_ and *TF*_4_ are significantly higher in NetREm (**Fig. S1D, Table S1:** p < 2e-16), highlighting NetREm’s grouped variable selection property that prioritizes them due to their strong corresponding PPIs with *TF*_1_ and *TF*_5_ that both strongly correlate with *y* (Li and Li 2008) (**Fig. S1E**). Unlike BRMs that favor *TF*_3_ over *TF*_4_ (since *TF*_3_ has greater |*r*| with *y*), NetREm prioritizes *TF*_4_ over *TF*_3_ since *TF*_4_ strongly interacts with *TF*_5_, while *TF*_3_ has weak PPIs with other TFs. Sensitivity analysis (**Fig. 2D-E**) for fixed *α* and *β* confirms |*c*^*^| for *TF*_2_ and *TF*_4_ increase as their respective PPIs with *TF*_1_ and *TF*_5_ strengthen. NetREm’s test MSE increases from 0.15 to 0.22 as *β* increases from 0.01 to 1 (**Fig. S1F**). Test MSEs for BRMs ≈0.14, which we can also achieve with NetREm(*β* = 0.01, LassoCV *α*); nonetheless, studies emphasize obtaining more interpretable and context-driven features by incorporating network information into models, even if accuracy is sacrificed (Li and Liu 2022). **Fig. S1G** illustrates changes in *B* as *β* increases from 0.01 to 2.

One of NetREm’s salient features is its ability to explicitly model and integrate PPIN structures/relations among TFs. By doing so, it identifies key TFs for predicting *y*, which may not exhibit strong *r* with *y* individually. NetREm also generates a coordination network among TFs that co-regulate TG, a capability beyond the scope of BRMs. Instead, BRMs, primarily focus on prediction accuracy by selecting *c*^*^ to best represent each TF’s impact on *y*, typically treating TFs as “lone wolves”, neglecting TF-TF interdependencies vital to GRNs. Our *B* network shows *TF*_1_ and *TF*_2_ cooperate with *TF*_3_ weakly to increase *y*, and *TF*_4_ and *TF*_5_ collaborate to decrease it; *TF*_1_-*TF*_2_ and *TF*_4_-*TF*_5_ links are highly-confident. NetREm predicts novel direct/indirect links like *TF*_1_-*TF*_5_ activator-repressor antagonism: *B* = -97.9. *TF*_3_ has weaker relations with *TF*_1_ than *TF*_2_ does (41.7 vs. 71.5) and the smallest |*c*^*^| = 3.4e-2.

We detail more simulations in **Fig. S2-5** and **Tables S2-3** in **Supp. Methods**. We show that results are stable and consistent for various expression sparsity levels and for *M* ≪ *N* cases as well, excess *β* over-constrains NetREm causing its predictions to suffer (bias-variance trade-offs), NetREm has more robust *c*^*^ estimates (less variable) and more accurate TF assignments, and it can capture complex TF-TF PPIs. Our toy data with a few TFs and 1 TG intuitively explains NetREm and its advantages.

Next, we benchmark NetREm in real-world settings: many TFs and TGs. We apply it step-by-step for each TG in a cell-type to learn *B*, TF-TG regulation, TF-TF coordination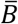.

### Benchmarking NetREm with No Prior GRN Info

Here, we assess how effectively NetREm predicts cell-type TF-TG and 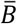 networks in real-world scenarios lacking prior GRN information (**Methods, Supp. File 5**). All TGs have the same *N* candidate TFs; for TGs that are TFs, *N* is reduced by 1. We anticipate that incorporating prior GRNs derived from multiomics data may enhance our performance by providing a tailored, context-specific list of promising TFs for each TG.

#### Evaluating TF-TG regulatory links

To evaluate our networks in terms of predicting regulatory TF-TG links as well as accurate TF roles (based on signed *c*^*^: +/-), we compare NetREm with BRMs for predicting signed-TF-TG links. We use SERGIO (Dibaeinia and Sinha 2020) to input a signed hESC GRN and generate 6 realistic datasets (1,250 TGs, *N* = 207), varying *M* and noise %. Overall, NetREm has the highest precision, indicating its superior reliability in correctly identifying true TF-TG links, assigning activator/repressor roles, minimizing FPs (**Fig. S6**).

We benchmark our inferred TF-TG links using single-cell expression data for human HSCs and mice (mESCs, mDCs) (**Figs. S7-8, Tables S4-6**) against gold standard GRNs (McCalla et al. 2023; Zhang et al. 2023). Due to its use of biological information (Shojaie and Michailidis 2009), NetREm demonstrates higher sensitivity in identifying relevant biomarkers, though it shows lower specificity compared to Lasso and ElasticNet (Li and Li 2008). By adjusting *β* and *α*, we can fine-tune NetREm’s behavior; increasing *α* might decrease sensitivity but enhance specificity. Our findings are consistent across updates in input PPINs, comparing well between mouse STRINGdb versions 11 (V11) and 12 in mESCs.

NetREm features a unique grouped variable selection mechanism that leverages PPIN structures to prioritize tightly-linked TF groups involved in known PPIs. This optimizes the selection of cell-type TFs that co-regulate TGs, enhancing NetREm’s ability to identify complex TF-TF relations. We highlight this in HSCs. ElasticNet and Lasso fail to identify critical TF-TG links, for instance, missing regulation of *ATF2*, a key HSC TF (Ju et al. 2023). Conversely, Linear Regression and Ridge predict that *ATF2* is regulated by all *N* = 177 TFs, showing their alarming potential for FPs. However, NetREm(*β* = 10) identifies 8 TFs for *ATF2*, with 7 confirmed by gold standards. This pattern extends to other TGs like *BRD2, RNF167, DUSP2*; NetREm identifies groups of both validated and novel TFs involved in biologically relevant PPIs for these TGs and more. For instance, 1 of 10 novel TFs for *RNF167* is TFAP4, which forms PPIs for adipogenesis and cell population proliferation regulation alongside 3 of 17 verified final TFs. NetREm effectively uses prior TF-TF PPIN knowledge to identify genuine TF-TF coordination for TG regulation, navigating through data noise and avoiding false correlations (*r*). It integrates *r* and PPIs among TFs, to achieve superior generalizability and consistency in identifying intricate TF-TF relationships that BRMs may struggle with.

Our TF-TG networks complement SOTA cell-type GRN tools. NetREm has comparable F1 Score, sensitivity, specificity, and accuracy with 10 tools benchmarked in mDCs (McCalla et al. 2023). No method outperforms others. However, NetREm uniquely infers *B* and cell-type 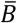, capabilities that other tools lack.

#### Evaluating TF-TF Coordination

We evaluate performance of our 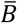 in mESCs, mDCs, and PBMCs using V11 PPIN as input (proxy for outdated information), categorizing TF-TF pairs into 4 groups based on their status in V11 and in updated V12: absent in both, present in both (TPs), removed in V12 (FPs), discovered in V12 (FNs). Top TF-TF links have higher 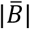. Welch 1-sided tests (p-adj < 0.05) compare 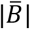 values across groups (**Tables S7-12, Figs. S9-11**). For instance, in mESCs, NetREm often reflects known PPIs, flags FPs to remove, and uncovers biological truths, nominating promising candidate PPIs for follow-ups. Overall, NetREm prioritizes known TPs and has potential to flag future TF-TF PPIs (FNs) that are currently unknown.

Further, we benchmark NetREm against the RTNduals (Chagas et al. 2019) tool. In 12 of 13 human contexts (**Fig. S12**), our top *k* links using V11 input PPIN outperform RTNduals, with a higher % that are verified PPIs in V12 and other sources. This underscores NetREm’s efficacy in identifying TPs and leveraging historical PPINs to predict previously unknown PPIs; this is encouraging as only a small number of ∼130-650k estimated human PPIs are currently known (Sevimoglu and Arga 2014; Venkatesan et al. 2009; Yu et al. 2020).

To validate NetREm’s capability to prioritize biologically relevant links, we leverage a Contextual PPI Database (CPPID) (Kotlyar et. al 2022) that annotates PPIs with >243 terms but lacks cell-type specificity (**Fig. S13**). By showing that our top links are indeed enriched for context-specific terms (e.g. NS-related in Microglia (Mic) and pooled SCs, immune-related in PBMCs), we suggest a potential extrapolation to cell-type specificity. This highlights NetREm’s pioneering potential to discover cell-type TF-TF PPIs, addressing a crucial gap in existing global PPINs.

#### Gene regulatory links between TFs and TGs in human mSCs & nmSCs

With emerging new single-cell epigenomic data from many human tissues, we can model GRNs in novel contexts. We apply NetREm to analyze SCs, which play a pivotal role in maintaining, regenerating, myelinating, and supporting PNS neurons. SCs are derived from neural crest and exhibit tremendous flexibility not only in myelination but also in several other tissues; they also function as terminally differentiated cells that can reverse differentiation after nerve injury to aid nerve regeneration (Ma and Svaren 2018). Single-cell rodent NS studies reveal substantial diversity in SC differentiation status (Gerber et al. 2021; Yim et. al 2022). 2 main SC phenotypes (or sub-types) are: 1) mSCs associated with larger diameter axons (>1 micron); 2) nmSCs that wrap a bundle of smaller diameter axons (typically sensory axons) i.e. a Remak bundle. While mSCs envelop axons in myelin sheaths to enhance conduction speed, nmSCs support sensory axon function/interactions without forming myelin, contributing to overall nerve integrity.

Uncovering TF-TG regulatory mechanisms modulating cellular processes in mSCs and nmSCs may help us understand and treat debilitating nerve injuries, demyelinating disorders, and hereditary neuropathies. Mutations affecting SC function are the most prevalent cause of demyelinating genetic neuropathy Charcot-Marie-Tooth disease (CMT) (Tao et. al 2019) and some affect major transcriptional TFs of SC differentiation like EGR2 and SOX10 (Srinivasan et al. 2012; Fröb and Wegner 2021). Both TFs are co-expressed in myelinating SCs, colocalizing at several myelin REs (Jones et al. 2007; Poitelon et. al 2016). Mutations in EGR2 may disrupt cooperative SOX10 TFBS binding (LeBlanc et. al 2006) and its subsequent regulation of TGs. SC regulation also involves TEAD1 and other Hippo regulators to govern shared TGs and orchestrate PNS myelination (Srinivasan et al. 2012; Lopez-Anido et al.

2016). Since there are no TFs that are exclusively expressed by SCs, uncovering TF-TF coordination networks crucial for SC lineage maturation and differentiation into mSCs or nmSCs also holds significance (Ma and Svaren 2018). However, distinct transcriptional regulatory networks coordinating SC function and underlying states and cell fates require a variety of TFs beyond EGR2 and SOX10 (Hung et al. 2015).

In response, we apply NetREm to each TG in mSCs and nmSCs using single-cell data for human Dorsal Root Ganglion (DRG). We derive prior GRNs using multiomics data (details: **Methods**). To do this, we annotate open chromatin regions in adult SCs with known RE peak-to-TG links (Zhang et al. 2021) and use motif-based analysis to predict TFs that may associate with these REs. By removing low-expressed TFs in corresponding training gene expression data for mSCs and nmSCs in the human DRG, we create tailored prior mSC and nmSC GRNs. For each TG in an SC-type, we input its *N* TG-specific selected candidate TFs (from respective prior GRN of initial TF-RE-TG links) to NetREm. In total, NetREm outputs 183,242 mSC and 277,541 nmSC total TF-TG links (**Supp. File 1**) comprising 221 TFs and 8,950 TGs in mSCs, 228 TFs and 5,207 TGs in nmSCs. Both share: 33,806 TF-TG links, 27,037 sign-TF-TG links, 3,841 TGs, 197 TFs. TF EGR2 is mSC-specific (Balakrishnan et al. 2021). We enhance our networks by overlaying them with prior GRN annotations, resulting in our finalized TF-RE-TG regulatory networks.

We examine NetREm’s results for 8 core SC TFs that have validation (genome binding and loss-of-function (LOF)) data from rodent SCs: EGR2, NR2F2, RXRG, SOX10, SREBF1, STAT1, TEAD1, YY1. LOF TGs, whose expression varies upon TF knockdown, show their direct or indirect dependency on the TF (Nie et al. 2020). Valid direct TGs are a subset of LOF TGs with ChIP-seq evidence of nearby TF binding, suggesting direct regulation by the TF (Titsias et al. 2012); this dual confirmation enhances confidence in these direct TGs (Badia-i-Mompel et al. 2023). RXRG lacks LOF data. In **Fig. 3A** we provide counts of NetREm-predicted: direct, all LOF, and novel candidate TGs. We also report on eTGs with strong Tibial Nerve eQTL support (THE GTEX CONSORTIUM 2020) based on mapped SNP-TF-RE-TG predictions: instances where an eSNP strongly alters TF binding to a RE, influencing TG expression (**Methods**). For example, we predict 2,015 YY1 TGs in nmSCs (139 direct, 304 LOF only, 1,572 novel), where LOF and direct TGs are significant (hypergeometric p < 0.05); of these, 40 direct (28.8%), 75 LOF only (24.7%), 488 novel YY1 (31%) TGs are eTGs. In both SCs, our final TF-TG predictions for all LOF and direct TGs have higher sensitivity and F1 scores compared to LassoCV and ElasticNetCV and higher accuracy and specificity than GRNBoost2, Linear Regression, RidgeCV (**Fig. S14**).

**Figure 3.**
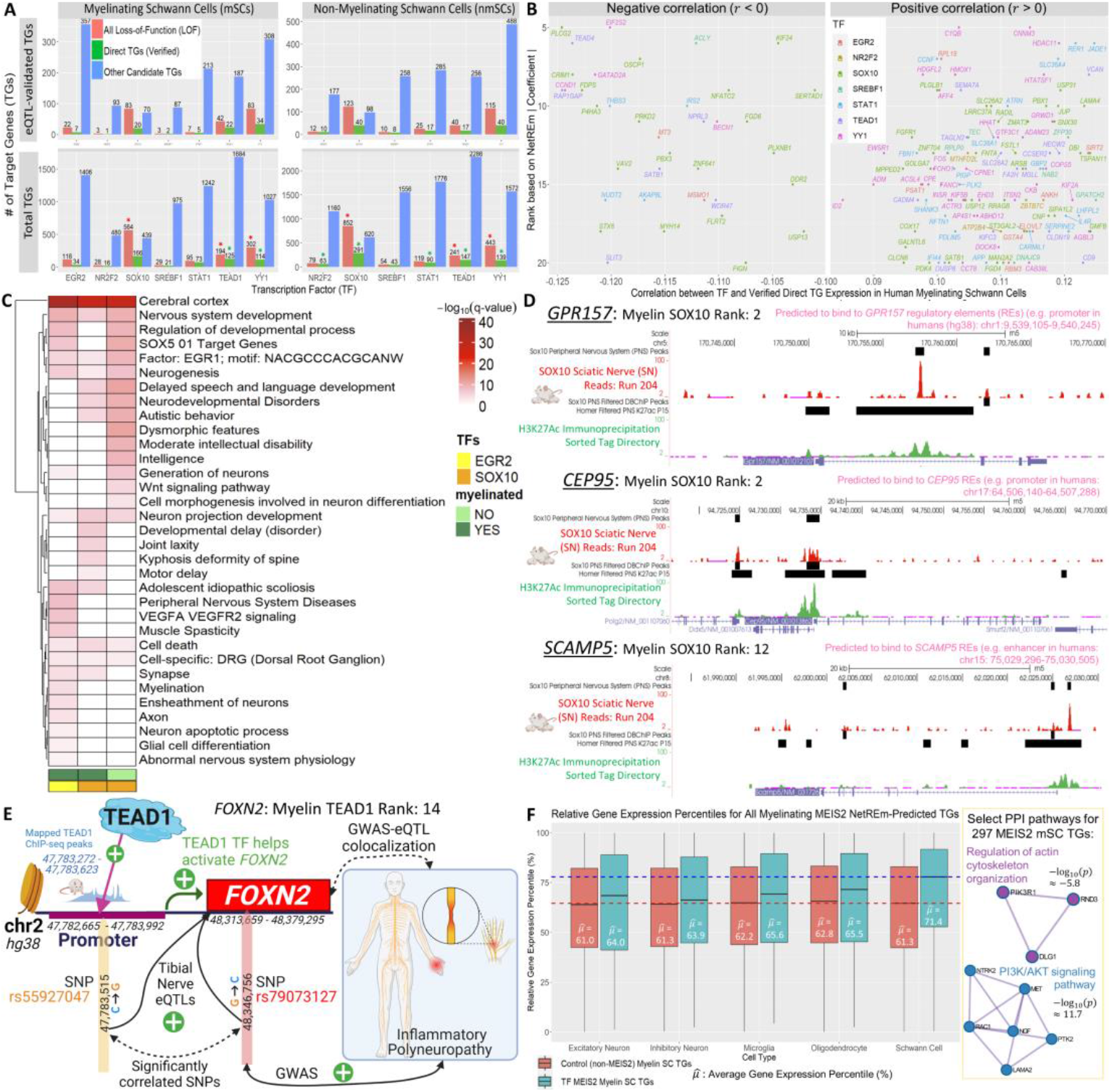
Gene regulatory links between TFs and TGs in human mSCs & nmSCs **A)** Bar plot categorizes TGs for core TFs in mSCs and nmSCs. Top: eTGs with eQTL validation. Bottom: original counts with star (*) for over-enriched TGs based on hypergeometric test (p-adj < 5%). **B)** Points are Direct TGs in mSCs. X-axis: *r* between core SC TF and direct TG in train data. Y-axis: rank of TF for regulating given TG, where smaller ranks imply greater |*c*^*^|, strong relations. For simplicity, we show results where the TF is the top 5 to 20 for its direct TG. **C)** Hierarchically clustered, select gene enrichments for all EGR2 mSC, SOX10 mSC, SOX10 nmSC TGs; values: − log_10_(*q*) **D)** Rat epigenome tracks for 3 novel mSC SOX10-activated TGs with SOX10 as a top predictor TF. Tracks: rat sciatic nerve ChIP-seq peaks, SOX10 PNS peaks, histone modifications associated with enhancers (H3K27ac immunoprecipitation sorted tags). Pink text: SOX10 binding in humans based on prior mSC GRN. **E)** Tibial Nerve eSNP rs55927047 (chromosome (chr) 2: 47,783,515 DNA change: C to G) in human *FOXN2* promoter (overlaps with orthologous TEAD1 rat ChIP-seq binding peaks) may strongly boost TEAD1 binding affinity to *FOXN2* promoter to activate (*c*^*^ > 0) *FOXN2*, a GWAS-eQTL colocalized biomarker for inflammatory polyneuropathy. *FOXN2* is a novel TEAD1 TG with support: TEAD1 rat ChIP-seq binding within 100 kilobases (kb) of its Transcription Start Site (TSS). **F)** Left: Boxplots compare relative expression percentiles for all MEIS2 mSC TGs in GTEx pooled SCs with those in 4 CNS cell types. Median percentile for MEIS2 mSC TGs in SCs overall is 71.4 versus ≤ 61.3 in controls (non-TGs for MEIS2 in mSCs). Right: enriched PPI paths for MEIS2 mSC TG proteins, which are important in SCs.

We explore cases where NetREm accurately identifies a core SC TF as the top predictor (low rank) for its direct TG, even when training expression data shows a low *r*(TF, TG) (**Fig. 3B**). This is important as studies observe that in eukaryotes (unlike in prokaryotes), correlations *r* and mutual information among TFs and known TGs are not much higher than those between TFs and non-TGs (Zaborowski and Walther 2020; Escorcia-Rodríguez et al. 2023). For example, SOX10 weakly correlates with *SIPA1L2* (Tao et. al 2019)(candidate modifier TG for CMT type 1A) but predicted to have rank 17. Despite its relatively weak *r* = -0.14 with *MBP*, a major constituent of myelin sheaths, STAT1 is its top 10 TF. Although *APP* exhibits a higher *r* with STAT1 compared to *FBN1* (9.7e-2 vs. 10.8e-2), NetREm ranks STAT1 higher as an activator for *FBN1* than for *APP* (13 vs. 20). This aligns with findings (Gu et. al 2022) of STAT1 LOF impacts of -0.61 for *FBN1*, -0.51 for *APP*.

NetREm reveals biologically relevant signals, identifying novel TGs for TFs. RXRG’s high regulatory activity in nmSCs (**Table S13**) is consistent with rodent/human studies (Gerber et al. 2021). All TGs across 3 groups (EGR2 mSC, SOX10 mSC and nmSC) are enriched in PNS-related terms (**Fig. 3C**). We find 2,316 SOX10 TGs (159 shared, 428 direct, 1,314 LOFs overall) with 29 direct and 103 LOFs overall in both SCs. **Fig. 3D** shows rat nerve epigenome tracks for 3 novel SOX10 candidate mSC TGs (*GPR157, CEP95, SCAMP5*) with strong SOX10 binding to SC regulatory regions in open chromatin (Lopez-Anido et. al 2015). In **Fig. 3E**, SNP rs55927047 enhances TEAD1 binding to its TFBS on *FOXN2*’s promoter to help activate *FOXN2* in adult mSCs. Orthologous rat nerve TEAD1 ChIP-seq peaks also overlap with this promoter. This eSNP correlates strongly (is in linkage disequilibrium (LD)) with rs79073127 that links to higher inflammatory polyneuropathy risk in Pan UK Biobank GWAS (Turley et al). *FOXN2* GWAS-eQTL colocalizes for this condition, with 75% probability (Wallace 2020). Both SNPs correlate with increased *FOXN2* expression. We highlight additional TFs in SCs, underscoring their significant roles despite limited validation. Notably, our 297 mSC TGs for MEIS2 (core TF in DRG sensory neurons (Roussel et al. 2022)) are prominent in PPI pathways like: PI3K-Akt signaling (crucial for PNS myelination (Ishii et al. 2021)), actin cytoskeleton organization (essential for PNS regeneration (Wang et al. 2018)) (**Fig. 3F**). These TGs have higher expression in pooled SCs compared to 4 CNS cell types (1-sided t, p-adj < 2e-6), a pattern absent in controls. **Fig. S15** showcases principal hub TFs like EGR1 (485 nmSC eTGs) and RXRA (971 eTGs), regulating the most eQTL-validated eTGs in SCs.

### Coordination among TFs for gene regulation in human mSCs & nmSCs

Our test MSEs are significantly lower than those of Linear and Ridge BRMs (**Fig. 4A, Table S14**). NetREm predicts SC-type-specific coordination *B* for each TG. It also outputs 22,809 mSC and 24,795 nmSC non-zero direct/indirect TF-TF coordination 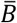 links (**Supp. File 2**). Notably, top CPPID contexts for strong mSC and/or nmSC 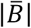 relate to DRG, brain (**Fig. S16A**). **Fig. 4B** shows 23 of 24 mSC-specific TFs in a mSC 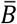 network of 77 known PPIs, excluding novel links (*w* = 0.01) for simplicity (nmSCs: **Fig. S16B**); POU3F1-EGR2 mSC cooperativity is extremely strong (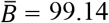 percentile), JUNB-ATF4 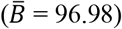 interact in PNS neoplasms like Schwannomas. BNC2’s regulatory activity in nmSCs may be attributed to the absence of its predicted repressor EGR2, ranked 10 of 20 mSC TFs for *BNC2*.

**Figure 4.**
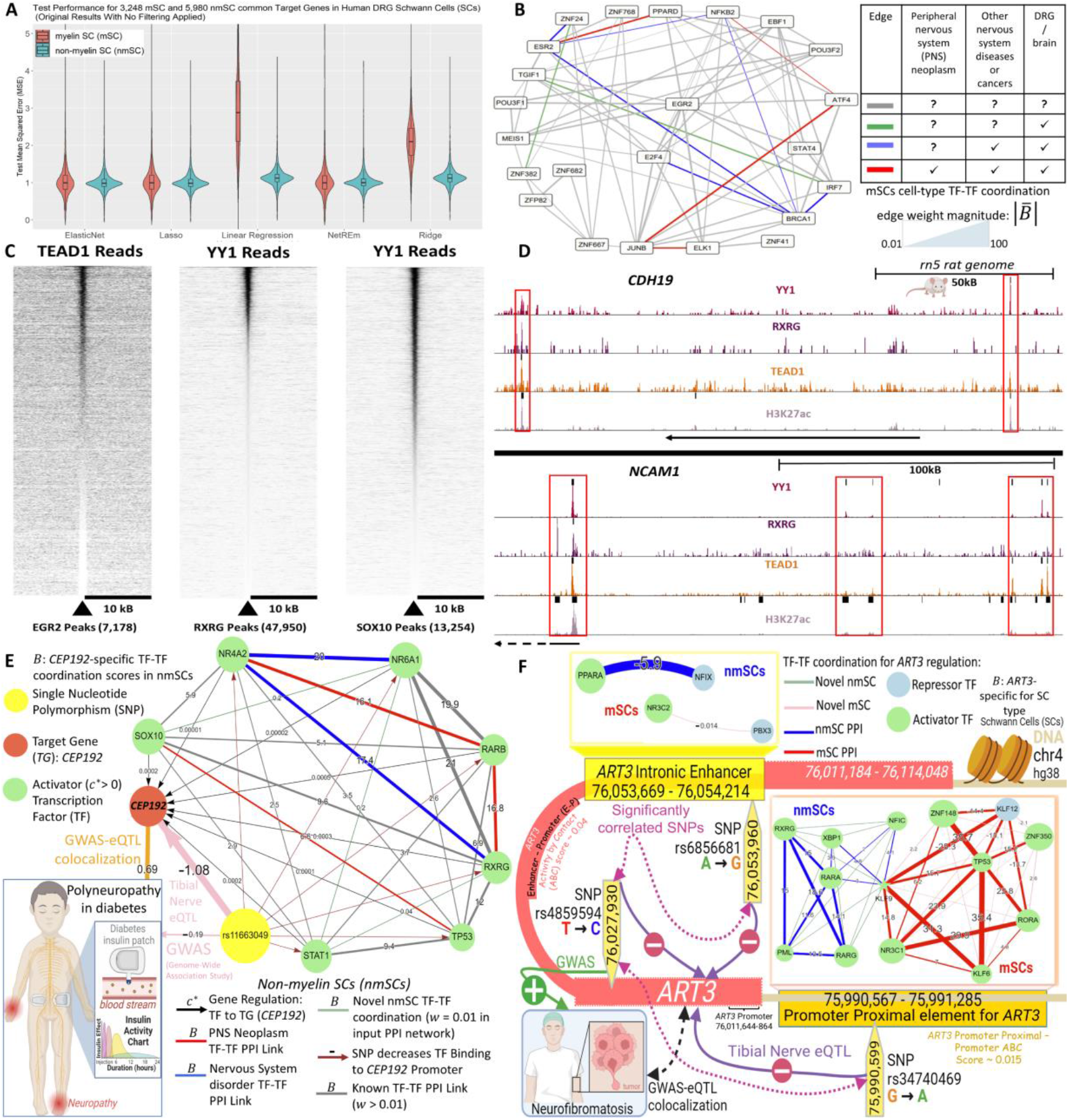
Coordination among TFs for gene regulation in human mSCs & nmSCs **A)** Density boxplots: NetREm outperforms Linear Regression and RidgeCV in both SCs, with lower test MSEs. NetREm predicts links for more TGs than ElasticNet and Lasso do. For mSCs, NetREm achieves median MSE: 0.95; nmSC: 1.1. **B)** Input PPIN subnetwork for only known TF-TF PPIs among 23 of 24 mSC-specific TFs only; artificial (novel) links excluded. Edges represent cell-type TF-TF cooperation 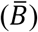 across TGs in mSCs since 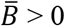 for all. High 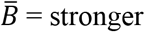 cooperation in mSCs. Edges annotated based on CPPID. **C)** C&R heatmaps show overlaps of TEAD1 centered on EGR2 peaks, YY1 C&R centered on RXRG peaks, YY1 C&R centered on SOX10 peaks. **D)** YY1, RXRG, and TEAD1 C&R tracks along with ChIP-seq H3K27ac tracks from rat S16 SC line. Boxes highlight enhancer regions where TFs colocalize across TGs *CDH19* (top tracks) and *NCAM1* (bottom tracks). Both TGs are impacted by Tibial Nerve eSNPs altering regulatory TF binding at their loci. **E)** *CEP192* in nmSCs GWAS-eQTL colocalizes with low risk of polyneuropathy in diabetes. Tibial Nerve eSNP (yellow) associates with low risk of this condition and low *CEP192* expression by potentially strongly disrupting bindings of 6 activators (green) to *CEP192* promoter. SOX10 cooperates with them. *CEP192*’s *B* links: novel (teal); known TF-TF PPIs colored based on CPPID annotations: PNS Neoplasms (red), NS disorders (blue), neither (grey). **F)** Tibial Nerve eSNPs potentially influence regulation of GWAS-eQTL colocalized neurofibromatosis TG *ART3* through activators and repressors forming distinct TRMs in mSCs and nmSCs along interacting REs (3D DNA loop). *ART3*’s (intronic) promoter has Activity by Contact (ABC) scores of ∼0.04 with intronic enhancer, ∼0.015 with proximal promoter RE. TF-TF *B* edge widths are shown relative to other TF-TF *B* links in that given *ART3* RE (i.e. intronic enhancer, promoter proximal).

NetREm discovers and prioritizes novel TF-TF coordination links that are promising. 48 of our novel links in nmSCs (37 also in mSCs), comprising 30 TFs, are validated by strong SAINT scores of physical TF-TF binding in recent BioID/AP-MS human experiments (Göös et al. 2022) (**Fig. S16C**). RXRG, TEAD1, and YY1 co-regulate 366 nmSC (**Fig. S16D**) and 15 mSC TGs, suggesting their preferential coordination in nmSCs. RXRG links with STAT1, TEAD1, and YY1 are unknown in our input PPIN (**Fig. S17A**); these 4 core SC TFs co-regulate 174 TGs in nmSCs, *SETD2* in mSCs. RXRG strongly positively correlates with TEAD1 and YY1 (*r* = 0.5, 0.7) in nmSCs, but negatively in mSCs (-0.9, -0.7) in mice sciatic nerves (Gerber et al. 2021). STAT1 has 591 (Jaccard Similarity (JS): 0.21), YY1 has 599 (JS: 0.20), TEAD1 has 843 (JS: 0.26, significant) co-regulated nmSC TGs with RXRG (**Fig. S17B-C**). In nmSCs, RXRG and YY1 share 94 eSNPs, 95 eTGs, 88 eQTLs compared with 28, 54, 24 for RXRG and STAT1 (cooperate for 704 TGs (**Fig. S17D**), antagonistic for 104 (**Fig. S17E**)); RXRG-cooperation 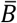is: TEAD1 (16.3), STAT1 (23.5), YY1 (24.1) (**Fig. S17F**).

To independently test our predicted TF-TF coordination for these core SC TFs, we use binding data from rat SCs in PNS, derived from ChIP-seq analysis of the active enhancer H3K27ac mark and ChIP-seq/Cut&Run read density assay (C&R) data of TF binding in nerve or S16 SC line. Of 15,864 ChIP-Seq H3K27 enhancer peaks shared between PNS and S16 lines, RXRG shares 3,450 and 2,017 peaks with YY1 and TEAD1 (**Fig. S17G**). 43.9% of RXRG peaks have YY1 binding (most colocalization among core SC TFs) and 25.7% of RXRG peaks have TEAD1 binding (**Fig. S17H**). Conversely, 24% of TEAD1 peaks and 28% of YY1 peaks have RXRG binding. C&R helps determine the extent of colocalized binding along REs. TEAD1 C&R centered across EGR2 peaks reveals TEAD1 colocalizes at ∼40% of EGR2 TFBSs (**Fig. 4C**), supporting predicted EGR2-TEAD1 coordination in mSCs. YY1 C&R reads overlap ∼70% when centered on RXRG peaks and SOX10 peaks. SC marker *CDH19* is preferentially expressed in nmSCs (Stratton et al. 2017) and has RXRG ChIP-seq binding nearby. Rs17799413 may associate with lower *CDH19* expression (slope: 2.4e-1), strongly altering binding of TEAD1 in both SCs and 7 core TFs (e.g. RXRG, YY1) in nmSCs. C&R sequencing reveals TEAD1 and RXRG binding peaks are 40 kb upstream of *CDH19*’s TSS (**Fig. 4D**). *NCAM1*, a YY1 LOF and SOX10 direct TG, codes for an adhesion molecule preferentially expressed in nmSCs (Martini and Schachner 1986). Our network predicts regulation by TEAD1, YY1, and RXRG in nmSCs. Rs10749999 associates with higher *NCAM1* expression (slope: 9e-2) and may boost YY1 and TEAD1 binding in nmSCs. Active SC enhancers ≈130 and 200 kb upstream of *NCAM1*’s TSS have TEAD1 and YY1 binding. *NCAM1* has TEAD1, RXRG, and YY1 binding at a promoter surrounding its gene locus. Rat H3K27Ac and TF ChIP-seq data shows RXRG co-regulates: 555 TGs with TEAD1, 352 with YY1, and 27 of 91 mapped STAT1 TGs. This further supports our predicted coordination by these 4 TFs, showing NetREm’s prowess in identifying novel colocalizing TFs in the absence of current evidence of direct binding interactions from high-throughput studies of PPIs.

Computational methods help decipher functional impacts of SNPs on PPIs, aiding in uncovering disease risk genes for targeted precision therapies. Most non-coding, disease-associated SNPs alter human PPIs rather than protein properties like folding or stability (Cheng et al. 2021). Integrating TF-TG regulatory (*c*^*^) and TF-TF coordination (*B*) networks with non-coding eSNP rs11663049 sheds insights on how PPIs associate with phenotype: polyneuropathy in diabetes (**Fig. 4E**). Dysregulated mitotic checkpoint regulators may lead to abnormal insulin signaling in diabetes (Choi et al. 2016). *CEP192* helps form mitotic spindles (Joukov et al. 2010) and colocalizes for this phenotype with 69% probability. The eSNP reduces the risk of this phenotype and *CEP192* expression and may strongly decrease binding of 6 activator TFs to *CEP192*’s promoter in nmSCs. *CEP192* is co-regulated by RXRG-STAT1 TG. RXRG-NR4A2, NR4A2-THRB associate with PNS neoplasms. SOX10, with its dynamic, cell-type-specific cooperation, works with these TFs; SOX family TFs achieve cell-type-specificity via partner TFs that facilitate TG regulation by binding to nearby SOX TFBSs (Stevanovic et al. 2021). SOX10-NFIA cooperate in nmSCs 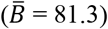 but display antagonism influencing glial lineage diversification (Glasgow et al. 2014) in mSCs 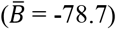.

To illustrate differences in TG-specific TF-TF coordination networks *B* between SC sub-types, even for shared TGs, we examine TG ART3 that colocalizes with neurofibromatosis (NF: characterized by the formation of NS-related tumors) with a 68% probability (**Fig. 4F**).We predict an interacting DNA chromatin loop (Zhang et al. 2021) featuring differing 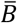 between mSCs and nmSCs involving strong repressors and activators (Sharov et al. 2022) to regulate *ART3*. 3 eSNPs link to lower *ART3* expression: NF-associated rs4859594 correlates with 2 regulatory SNPs (rs6856681: intronic enhancer; rs34740469: promoter proximal RE) that may disrupt coordination networks by strongly decreasing activator binding to *ART3’s* REs, increasing it for repressors. In both SC types, common KLF9 coordinates with TFs at *ART3’s* promoter, eagerly cooperating with: XBP1 (*B* = 4.84) in nmSCs, KLF6 (*B* = 31.3) in mSCs. PPARA-NFIX (nmSCs) and NR3C2-PBX3 (mSCs) are antagonistic relations on the enhancer.

### Prediction and comparative analysis of cell-type coordination among TFs for gene regulation across neuronal and glial cell types in Alzheimer’s disease (AD)

Cell-type-specific TF-TF coordination networks 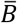 are crucial for neuronal functions like synapse plasticity and neurotransmission, and are disrupted in AD, leading to memory loss, neuroinflammation, cognitive decline (Mathew et al. 2022). Understanding how altered 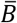 impacts TG expression in AD is essential for identifying master regulators and developing targeted therapies (Wang et al. 2016). Signaling PPIs associated with dementia symptoms highlight the potential of targeting these altered PPIs (Mao et al. 2020) to delay AD progression (Vargas et al. 2018).

In response, we integrate multi-omics data to construct 16 context-specific prior GRNs for 8 cell types in AD and Controls. For each TG in a context, NetREm uses the *N* TG-specific candidate TFs from the respective prior GRN as input features, based on TFs that may associate with its REs to regulate it (**Methods**). Ultimately, NetREm generates 16 corresponding TF-RE-TG (TF-TG links: **Supp. File 3**) and 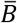 (**Supp. File 4**, word cloud: **Fig. S18A**) networks. Reverse engineering changes in networks across cell types may help illuminate molecular drivers of AD.

We explore TG-specific coordination *B* in Control vs. AD stages for 2 AD risk genes (Jia et al. 2020; Bossaerts et al. 2022) (**Fig. 5A**). *TMPRSS15* in Mic (*t*: 36.1, p-adj < 4.8e-283) and *ABCB5* in Inhibitory Neurons (InNs) (*t*: 22.6, p-adj < 4.5e-112) show notable increases in *B* in AD versus baseline Controls for the respective cell-type. Some TF-TF pairs strongly cooperate (i.e. 50 ≤ *B* ≤100) exclusively in AD: ZBTB14-ZNF281, FLI1-TAL1 for *TMPRSS15*; ZNF331-ZNF354A, MYEF2-SOX2 for *ABCB5*. For *TMPRSS15*, known AD links IRF7-STAT3 and STAT3-STAT5B have strong antagonism (i.e. -100 ≤ *B* ≤ -50) in controls but cooperate in AD. **Fig. 5B** compares strong 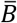 among select TFs between conditions for Mic and InNs. IRF7-STAT3 is in Mic, MYEF2-SOX2 is in InNs only. AD link STAT3-STAT4 is antagonistic in Controls for *TMPRSS15* in Mic and in Control Mic overall (but cooperative for other 3 networks). RORA-ELK1 is cooperative in all 3 but antagonistic in Control Mic. FOSL2-BACH1 is antagonistic in Mic but cooperative in InNs. Indeed, RORA activity increases in InNs and Mic in AD (Acquaah-Mensah et al. 2015). AD-annotated links in AD InNs/Mic include ELK1-SPI1 and ELK1-STAT3.

**Figure 5.**
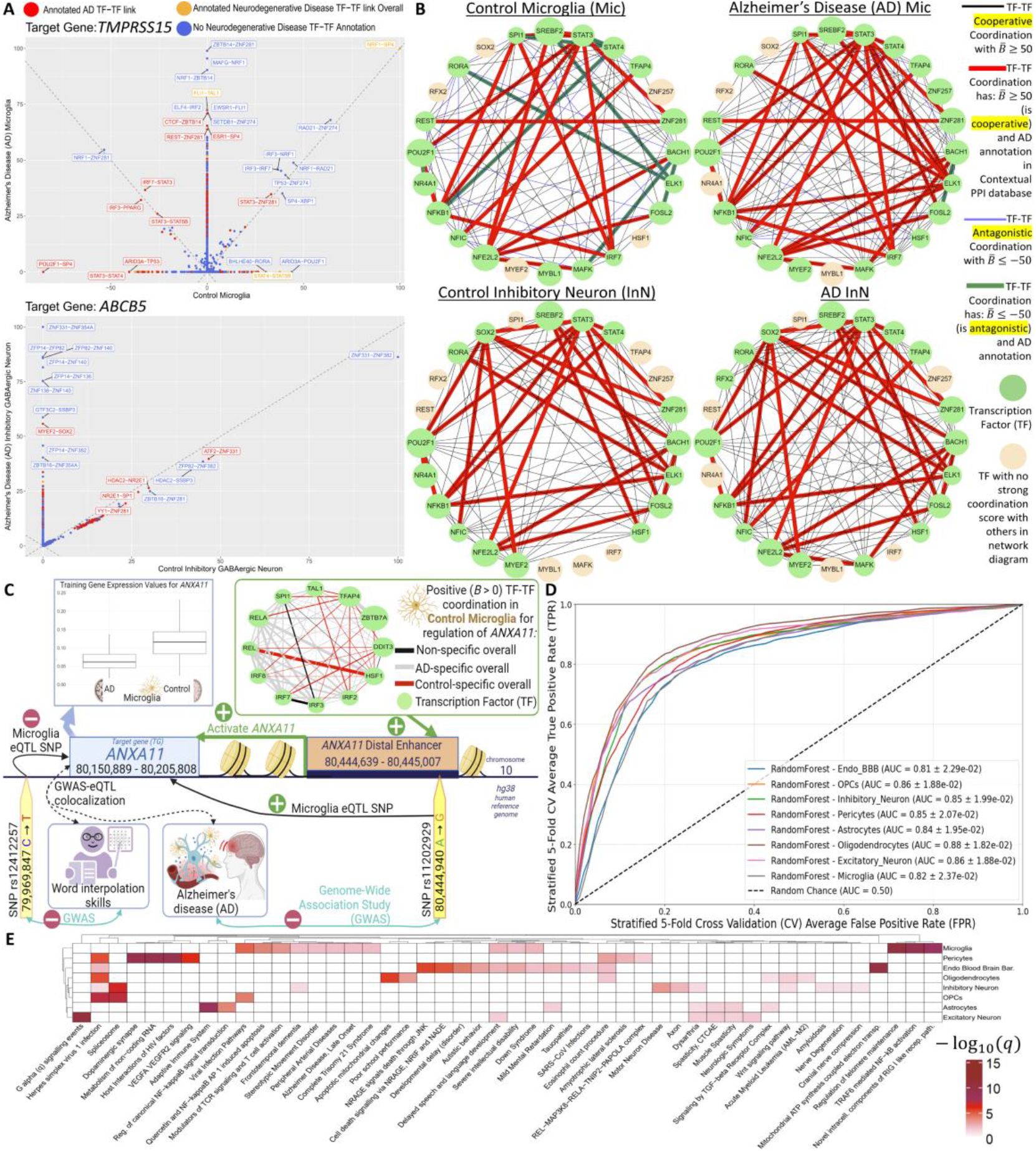
Prediction and comparative analysis of cell-type coordination among TFs for TG regulation across neuronal and glial cell types in AD. **A)** Coordination scores *B* for 2 AD risk TGs, *TMPRSS15* for Mic and *ABCB5* for InNs, in Control versus AD. Red and orange points are TF-TF links annotated by CPPID for AD and other neurodegenerative diseases, respectively. Blue points: no annotation for neurodegeneration. **B)** Circular network diagrams visualize strong cell-type direct/indirect TF-TF coordination in AD/Control Mic and InNs, focusing on light green TFs that exhibit potentially strong: cooperation 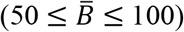 or antagonism 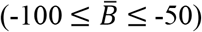. Thick edges are AD-related in CPPID. Peach: TFs not expressed or lacking strong links with other TFs in this visual. **C)** 12 TFs that may cooperate to activate *ANXA11* in TF-TG regulatory network in Control Mic (higher expression for *ANXA11* than in AD). Widths for TF-TF links are *B* scores for *ANXA11* regulation in Controls and colors are based on statistical significance of TF-TF links across all TGs in both Mic networks. Light gray links: higher in AD; red: higher in Controls; black: not significant. A SNP correlating with lower AD susceptibility, increases binding of this TF-TF network, and links to higher *ANXA11* expression. Risk SNP for reduced cognitive ability (poor word interpolation skills) links to lower *ANXA11* expression. **D)** Average stratified 5-fold CV ROC curves for RF models predicting TF-TF links annotated in neurodegenerative diseases (class 1) or not (class 0) for balanced classes. **E)** Enrichment analysis for top 500 optimal genes identified by RF for each of 8 cell types. Hierarchical clustering is performed on rows (cell types) and columns (terms), and − log_10_(*q*) values are shown.

**Fig. 5C** presents a multifaceted network weaving together TF-TG regulatory links, TF-TF coordination, phenotypes, SNPs. TF-TF coordination is discernibly stronger and positive in Control Mic (1-sided *t*), pointing to potentially disrupted cooperativity during AD. Our attention is drawn to regulation of *ANXA11*, a critical player in diverse functions (e.g. apoptosis, neutrophil function) and signaling paths (e.g. MAPK, P53) (Mirsaeidi et al. 2016). Mutations in *ANXA11* are correlated with NS diseases (Wang et al. 2022) and high risk of inflammatory conditions like sarcoidosis (Smith et al. 2017). Our AD-Covid study (Khullar and Wang 2023) assigns *ANXA11* to a Hippocampal Control gene co-expression module. NetREm offers nuanced insights on non-coding SNPs for AD (rs11202929) and cognitive function (rs12412257), linked to more and less *ANXA11* expression, respectively, in Mic (resident CNS macrophage immune cells). Rs12412257 associates with reduced word interpolation ability, a measure of fluid intelligence and reasoning (Turley et al) and prognostic marker of AD (Eyigoz et al. 2020). Rs11202929 is protective against AD (GWAS slope < 0) and may enhance binding affinity of 12 cooperating activators to TFBSs on *ANXA11’s* enhancer in open chromatin in adult Mic. SPI1 is a core Mic TF for AD genes (Rustenhoven et al. 2018). *ANXA11* has higher Mic expression in controls than in AD (*t* p-adj = 2.6e-68, log_2_(Fold Change) of (Control/AD) = 0.78; 668 AD, 676 controls). Our significant GWAS-eQTL colocalization analyses reveals that *ANXA11* expression positively associates with lower AD risk and better word interpolation ability. NetREm provides a powerful framework to deepen our understanding of complex GRNs and their implications across a spectrum of health conditions.

We further evaluate our 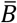. **Fig. S18B** compares 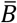 across 16 networks for 216 novel links validated in recent physical human experiments (Göös et al. 2022). In control ExNs and InNs, *r* ≈ 0.54 and 0.45, respectively, between 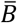 and JS of ChIP-seq overlaps (proxy for cooperativity (Yu et al. 2015)) for 6 TFs in neural cells (**Fig. S19**).

For each cell-type, we build default Random Forest (RF) (**Fig. 5D)**, Logistic Regression, Naïve Bayes, and XGBoost models (Pedregosa et al. 2011) to detect TGs with altered *B* from Controls to AD that may predict TF-TF links annotated (CPPID) with neurodegenerative disease (**Fig. S20**). Input data consists of changes in *B* across TGs from Control to AD, for the given cell type. To tackle this positive unlabeled learning problem (Yang et al. 2014), we undersample class 0 so each cell-type has an equal count of TF-TF links in both classes (**Supp. File 5**). We evaluate performance via stratified 5-fold CV noting RF is the most optimal approach as all 8 RF cell-type models have average area under Receiver-Operator Characteristic (ROC) curve (AUC) ≥ 0.81. Across our models, the top 500 RF feature TGs are enriched for neurodegeneration, cell-type functions, immunity, intellectual disabilities, tauopathy (**Fig. 5E**). OPCs and InNs have the highest overlap (35 top TGs); disrupted InN signaling to OPCs may diminish myelination and CNS interneuron activity, and severely impair prefrontal cortical network functions and social cognitive behavior (Fang et al. 2022) (**Table S15**).

## Discussion

In this paper, we present NetREm, a computational multi-omics-based approach that employs network-regularized regression on single-cell gene expression data to predict cell-type coordination among TFs for TG regulation. NetREm addresses a major challenge in traditional studies of cell-type GRNs. Gene expression data, often nascent, sparse and noisy, fails to capture crucial GRN mechanisms such as TF binding to DNA, coordination among TFs/cofactors, and DNA accessibility, and typically provides weak signals for distinguishing TP from FP TF-TG links (Kim et al. 2023; Badia-i-Mompel et al. 2023). Sole reliance on expression data for GRN inference is therefore woefully inadequate, and perhaps even futile, often leading to unstable and inaccurate results.

Explicitly modeling direct and indirect TF-TF interactions can enhance GRN inference, enabling discovery of novel TF-TG links and key cell-type TFs (Skok Gibbs et al. 2022). Functionally-related TF predictors, like neighbors in scale-free feature networks (e.g. TF-TF PPINs), can coordinate synergistically or antagonistically in biological processes like TG regulation (Kong and Yu 2018). Nonetheless, traditional methods often miss such complex dynamics of TF-TF PPIs involved in GRNs (Yazaki et al. 2016).

SOTA GRN tools like SCENIC indirectly hint at TF-TF interactions by analyzing TFs that co-regulate multiple TGs. However, these tools primarily focus on TFs with strong motif binding, excluding other prior information. For instance, in our comparative analysis, SCENIC identifies many TFs in SCs (nmSCs: 640, mSCs: 522) but overlooks TEAD1 in both SCs, instead detecting 3 other TEAD family TFs. TFs like TEAD1, which exhibit relatively weak motif-binding signals, are drowned out. In contrast, by incorporating TF-TF PPINs, NetREm effectively captures essential GRN relations for core SC TFs like TEAD1. This underscores the importance of integrating comprehensive prior information like PPINs, in GRN predictions from expression regression, a capability that NetREm successfully implements (Dibaeinia and Sinha 2020).

NetREm reveals cell-type coordination among TFs, 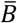, with some mediated by physical and others by indirect (e.g. pioneer/settler models show TFBSs are often >50 bp apart in REs) PPIs (Martin et al. 2023). By weighing known direct/indirect PPIs in the context of TG regulation, NetREm helps characterize existing PPINs at a cell-type level (Johnson et al. 2021; Murtaza et al. 2022; Hsu et al. 2022). It also helps address the link prediction problem, flagging undiscovered PPIs for follow-ups (Singh and Vig 2017). The lack of cell-type PPI annotations, while a challenge, offers NetREm an opportunity to contribute to ongoing efforts to broaden understanding of PPIs and protein dynamics with its dual capacity to annotate known TF-TF PPIs and discover novel cell-type-specific ones (Yu et al. 2023).

NetREm predicts unprecedented, weighted, cell-type-specific TF-TF coordination networks 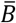 across various conditions, including both human and mouse contexts, even in the absence of prior GRNs. Our benchmarking shows our TF-TG regulatory networks not only perform competitively with SOTA GRNs but also uncover novel cell-type TFs that coregulate TGs; further, our 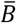 effectively prioritizes TP and FN TF-TF links.

Disrupted cell-type PPIs are critical in neurobiological disorders, since PPIs mediate neuronal functions (Mathew et al. 2022). We integrate multi-omics data, capturing various levels of TG regulation (e.g., scRNA-seq, scATAC-seq), to learn prior GRNs for NS cell-types for our SC and AD human applications. Detecting these candidate GRN TFs for TGs is key to inferring biologically significant cell-type TF-TG links (Zaborowski and Walther 2020; Zhang et al. 2023). Aligning NetREm’s TF-TG links with prior GRNs helps us deduce TF-RE-TG links. NetREm uncovers novel TF-TF crosstalk for TG regulation in SCs and during AD in neurons/glia.

Insights derived from NetREm may contribute to advancing targeted therapies and regenerative medicine. We apply our predicted regulatory and coordination networks to trace how non-coding eSNPs may alter co-regulatory dynamics among TFs, potentially altering expression of disease-associated eTGs.

NetREm extends to any discipline where predictors exhibit a network structure that informs the outcome. It expands upon previous work in network regularized regression by learning and generating embeddings using SVD. In the future, we can incorporate nonlinear dimensionality reduction into NetREm to capture nonlinear patterns and regularize latent representations with prior information. We may see if any final TFs form homodimers or adapt NetREm to account for this. Additionally, regularization networks can integrate other information, like signaling pathways, to learn TF-TF coordination for TG regulation. Beyond expression regression, we can extend NetREm to other emerging single-cell omics like scATAC-seq to explore TF and chromatin interactions in open regions.

## Methods

### NetREm Methodology

#### Integrating multimodal data & networks in NetREm workflow

##### Preliminary definitions

Proteins (e.g. TFs), and TGs (*italics*) are represented by HGNC symbols. TF gene expression is a proxy for TF protein abundance, assuming high gene expression (quantifies mRNA abundance) translates to high protein expression. NetREm can be applied to both single-cell and bulk expression data (sample by gene). Bulk data represents pooled collections of cell lines or tissues, yielding averaged expression profiles, as seen in patients.

We start with single-cell gene expression data for *M* samples (individual cells) and 𝒯 genes in a cell-type; scRNA-seq data is typically high-dimensional (*M* ≪ 𝒯), sparse, non-negative. We focus on *G* TGs, (i.e.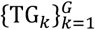), with expression profiles 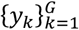, respectively, and 𝒩 potential cell-type TFs, where *G* ≤ 𝒯, 𝒩 < 𝒯. If all TFs are master regulators: 𝒯 = *G* + 𝒩; else (some TGs are TFs): 𝒯 < *G* + 𝒩. We ensure a TG is not its own candidate TF.

NetREm can use prior GRN information that identifies 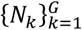 respective promising candidate TFs for the set of *G* TGs where *N*_*k*_ ≤ 𝒩 and varies based on the optimal prior GRN TFs selected for the given TG_*k*_. When prior GRN information is absent, TG_*k*_ (and other *G* − 1 TGs) will have the same fixed *N*_*k*_ = 𝒩 candidate TFs; it is to be understood that if TG_*k*_ is also a TF, its self-TF is excluded, so it has *N*_*k*_ = 𝒩– 1 candidate TFs.

These steps below are repeated for each of the *G* TGs. For simplicity, we explain the pipeline for predicting expression of a single TG. We use *N* to represent the # of its candidate TFs for this TG, and *y* for its true expression.

##### Optional (recommended) prior cell-type GRN information

Constructing candidate links from TFs to TGs may improve the quality of NetREm’s solution via initial feature selection of *N* biologically meaningful TFs tailored for TG, where *N* < 𝒩. Using diverse information on expression regulation guides the regression (Zaborowski and Walther 2020). We integrate prior GRNs (initial TF-RE-TG links) from multi-omics data in applications 6-7. When no prior GRN is used: if TG is a TF, it has *N* = 𝒩 − 1 candidate TFs, otherwise it has all 𝒩 TFs as candidates.

##### PPI network (PPIN, network prior)

Our comprehensive, weighted, undirected PPIN 𝕎 illuminates biological interactions among proteins (network nodes) with strong functional association evidence in the organism. It inherently captures direct (e.g. complex formation, transient interactions) and/or indirect (e.g. participate in shared processes or bind by intermediate hidden partners (De Las Rivas and Fontanillo 2012)) PPIs. Alas, PPINs attribute weight *w* > 0 to all PPIs, even to those with antagonistic, competing roles in paths (Szklarczyk et al. 2023); each edge *w* accounts for uncertainty in 𝕎 and is proportional to the probability that the 2 connected nodes interact (i.e. their integrative functional essentiality in 𝕎) (Li and Liu 2022). For the TG, we subset 𝕎 to obtain *W*^(0)^ that captures known TF-TF PPIs, reflects potential structure/relation background information among TG’s *N* candidate TF proteins, and is symmetric (i.e.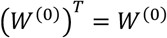. Higher *w*_*ij*_ denotes more confidence that *TF*_*i*_ and *TF*_*j*_ partner, directly and/or indirectly, in processes like regulating DNA chromatin loops of interacting REs for TG regulation (Wang et al. 2021). Here, *w*_*min*_ = m*i*n{*w*_*ij*_ ∈ *W*^(0)^} is the smallest confidence for known direct/indirect TF-TF links among *N* TFs. To enable NetREm to consider candidate TFs missing from *W*^(0)^, we add artificial weight 0 < η < *w*_*min*_ for missing pairwise edges; these novel edges may not exist (i.e. TFs truly do not coordinate: True Negatives) or are yet to be discovered (FNs). We obtain our final, fully-connected, TG-specific, TF-TF input PPIN: *W* ∈ ℝ^*N* × *N*^. *W* has 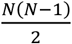 un*i*tless, global PPIs with *w*_*ij*_ > 0 equal to: 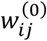 for known and η for artificial TF-TF links. We do not consider self-loops; instead, we set 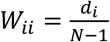 where *TF*_*i*_’s degree (connectivity) with other *N* −1 TFs is *d*_*i*_ = Σ_*k*≠*i*_ *w*_*ik*_ > 0.

##### NetREm integrates input data

For the *M* samples, the *X* ∈ ℝ^*M* × *N*^ matrix contains expression data for *N* predictors, while *y*^(0)^ ∈ ℝ^*M*^ is the expression vector for TG. We standardize each column of 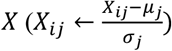 and 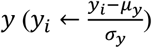 by respective means *μ*_*j*_, *μ*_*y*_ and standard deviations *σ*_*i*_, *σ*_*y*_, making them unitless. Then, each TF in *X* has 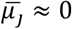 and *σ*_*j*_ ≈ 1; *μ*_*y*_ ≈ 0, *σ*_*y*_ ≈ 1. Pairwise *r* are preserved. If *M* < *N, X* suffers the curse of dimensionality. With technological advances and the advent of large single-cell sequencing studies, we anticipate a boost in *M* so *M* ≫ *N* will soon be the norm (Cuomo et al. 2023), especially since *N* ≪ 𝒯 (relatively few genes are transcribed and translated to proteins that are TFs). NetREm identifies which of *N* TFs can predict TG expression *y*, considering PPIs among TFs. NetREm can comprehensively integrate multi-omics data and PPINs to discover key TFs and TF-TF coordination events for TG regulation in a cell-type-specific manner.

## Step 1: Network regularized regression (Problem Definition)

Given gene expression data with *M* samples (rows) and *N* features (columns) represented as *X* ∈ ℝ^*M* × *N*^ (e.g. *N* candidate TF expression values for each *M*) and response *y* ∈ ℝ^*M*^ (e.g. TG expression for each *M*), we want to learn a linear predictor *y* ≈ *Xc*^*^ for *TG*. By imposing prior TF-TF PPIN information as a regularization term, we develop a functional map highlighting connectivity patterns among *N* TFs for TG; this typically steers NetREm to favor groups of TFs with shared PPIN connectivity over isolated TFs, thereby enhancing its ability to capture biologically relevant PPIs (Li and Liu 2022). Coefficients *c*^*^ ∈ ℝ^*N*^ represent the importance of TFs for regulating TG and are found by optimizing the following problem with objective function:

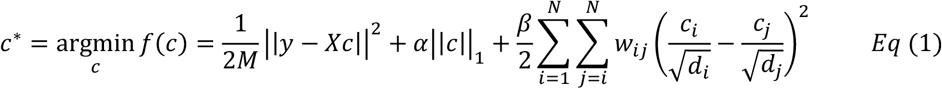

Our 3 terms in equation *Eq* (1) are unitless and compatible for addition: (data-fitting): ensures *Xc* is close to *y*, and 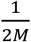 is a normalization factor to make it invariant to sample size *M*. **2**(sparsity-prior): favors a sparse solution (small # of non-zero *c*^*^), helping simplify the model and boost reliability. **3** (network-prior): penalizes differences between *c*^*^ of connected TF nodes, normalized by their respective network centrality *d* and adjusted for their global (organism-based) PPI weights with other candidate TFs in TG-specific input *W*. Inspired by (Li and Li 2008), this approach allows for a more equitable representation of TFs, irrespective of their *d*; this network-oriented variant of Ridge *L*_2_ penalty 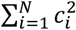, promotes topology-aware *c*^*^ shrinkage and smoothing for neighboring *TF*_*i*_ and *TF*_*j*_, with probability proportional to *w*_*ij*_. It underscores the principle that strongly connected TFs likely perform shared functions, even if their influence on TG expression (*c*^*^ signs) differs. This recognizes the community structure in existing PPINs that groups proteins with similar biological roles with *w* > 0, not distinguishing between cooperative (+) and antagonistic (-) PPIs (Padi and Quackenbush 2015; Szklarczyk et al. 2023). NetREm leverages this refined understanding, offering a comprehensive perspective on the interplay of TFs in *W*.

We can tune 2 hyperparameter knobs: network-constrained prior, *β* > 0, decides strength of PPIN regularization penalty (applied 1^st^: higher *β* guides NetREm to prioritize TFs with strong PPIs); *α* ≥ 0 impacts ℒ_1_ penalty (applied 2^nd^). NetREm, a PPIN-aware adaptation of ElasticNet, performs automatic variable selection based on expression and PPIN data, grouping and selecting strongly-connected TFs (emphasizing known TF-TF PPI subnetworks) in a spirit akin to ElasticNet. If *M* ≪ *N*, ElasticNet and NetREm may still select ≤ *N* TFs as final; this addresses limitations of Lasso regression that may indiscriminately select only 1 TF from a group of highly correlated TFs and only ≤ *M* TFs *i*f *M* ≪ *N* (Zou and Hastie 2005; Li and Li 2008).

## Step 2: Gene expression embeddings from network regression

Our novel method transforms the original problem into a Lasso regression problem in a new space with cell-type-specific TF-TF interactions. 1^st^, we represent the network-prior term in a more compact matrix-vector form as:

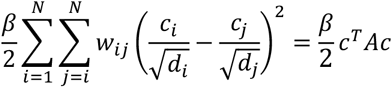

where *A* ∈ ℝ^*N* × *N*^, a symmetric normalized Laplacian matrix, is computed as:

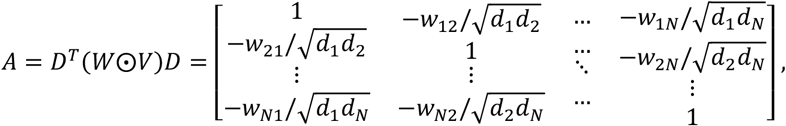

where 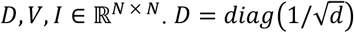 is a diagonal matrix with elements 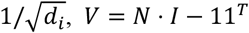 is a constant matrix where 1 ∈ ℝ^*N* × 1^ is an all 1 column vector and *I* is an identity matrix. *W⨀V* is element wise multiplication of *W* and *V*. Using this matrix-vector representation, we reformulate *f*(*c*) in *Eq* (1) as

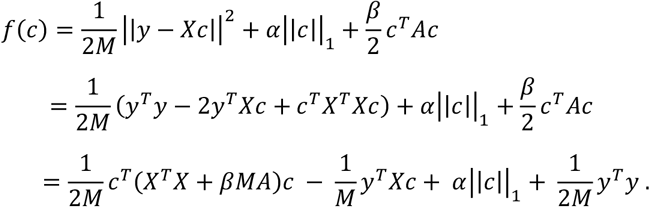

We set 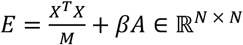, which is symmetric and positive semi-definite (psd) since *A* and Gram matrix *X*^*T*^*X* ∈ ℝ^*N* × *N*^ are symmetric and psd and *β, M, N* > 0. Cell-type-specific data term 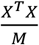 represents *r* among TFs in the expression data, the covariance matrix of *X* scaled by 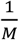. Cell-type-independent PPI term *βA* is based on global PPI *W* and is *β* for main-diagonals and is −*β* ≤ *βA*_ij_ < 0 for *i* ≠ *j*. Thus, 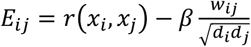, which balances *r* with PPIN knowledge (for a given *β* value) and information 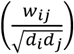.

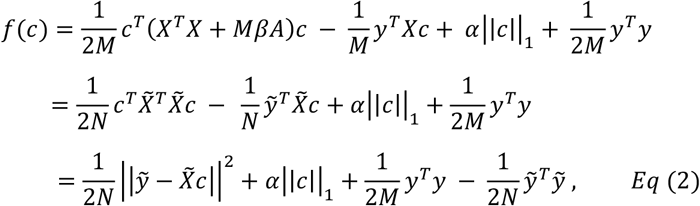

where 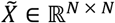 and is symmetric (i.e.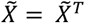) and 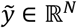 satisfies:

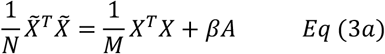

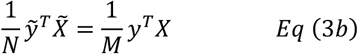

Finally, we reformulate *Eq* (2) as a conventional Lasso problem (by omitting constant term 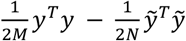 that we solve using existing standard Lasso or LassoCV (Pedregosa et al. 2011) regression solvers:

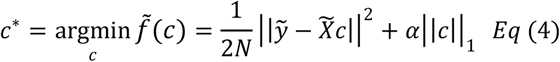

To compute 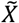 and 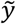 we perform SVD on *E* expressed as: *E* = *U*∑*U*^*T*^ where *U*, ∑ ∈ ℝ^*N* × *N*^. Here, *U* has left singular vectors and ∑ is a diagonal matrix of non-negative singular values 𝒮 = {*s*_1_, *s*_2_, …, *s*_*N*_} of *E*. Then, 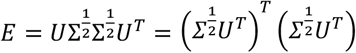. Based on *Eq* (3*a*),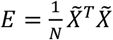. Then,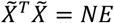. Hence,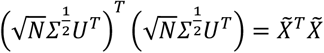. For improved stability, we set small singular values (e.g., *s*_*i*_ < 10^−6^*s*_max_) to 0, resulting in truncated ∑_*tr*u*nc*_, and similarly, we adjust inverse ∑_trunc_^−1^ setting inverse elements corresponding to small singular values to 0. By substituting 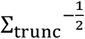 in place of 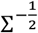, we effectively use truncated SVD, enhancing NetREm’s robustness by excluding contributions from small singular values. Thus, *X* and *y* are transformed to a new latent space of gene expression embeddings that incorporate PPIN information: 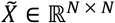 and 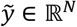, respectively, in *Eq* (4):

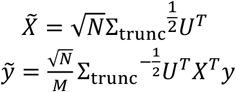

When *β* = 0, the transformation yields *X*’s principal components (PCs) via SVD on *X*^*T*^*X*, a trivial case without PPIN information; however, since we require 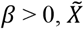 not only reflects its PCs but also includes PPIN structure, with *βA* added to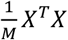, creating an “embedding”. This comprehensive approach captures both data patterns and PPI relations. We perform Lasso regression to solve *Eq* (4) for 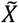 and 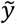, determining optimal *c*^*^. **Supp. File 5** provides more details.

## Output 1: Identification of novel cell-type TFs for regulating TG

### TF-TG regulatory network for TG

Several TFs regulate transcriptional activity of TG in a cell-type at a certain time (Nie et al. 2011). Solving the network-regularized regression problem produces a vector of Lasso *c*^*^ ∈ ℝ^*N*^ for *N* TFs predicting true TG expression *y*. We focus on *c*^*^ ≠0, which represents *N*^*^ final TFs for TG out of *N* candidates, where 0 < *N*^*^ ≤ *N*. NetREm constructs a comprehensive directed TF-TG regulatory network (complementary GRN) of *N*^*^ edges, weighted by *c*^*^. Here 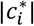, indicates the strength of the *TF*_*i*_-TG link, measuring *TF*_*i*_’s relative importance in regulating TG. *TF*_*i*_ with 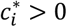 may activate TG and *TF*_*j*_ with 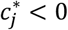 may repress TG transcription and subsequent expression. This tug-of-war between activators and repressors orchestrates TG regulation. Biological complexity enables certain TFs to have dual-function roles, alternating between activation and repression depending on context and signals (Skok Gibbs et al. 2022; Boyle and Després 2010). Given the competitive nature of TFBS binding, *N* − *N*^*^ discarded TFs may lose to some of the *N*^*^ TFs (i.e. antagonistic relation), but we do not speculate on this. Overall, NetREm unearths novel cell-type-specific, coordinating TFs involved in TG regulation, providing a more nuanced view of TF-TG interactions.

We evaluate our performance in training and testing expression data, comparing predicted 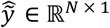 to actual 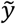 using metrics like Mean Square Error 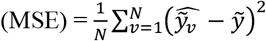. To achieve more accurate regulatory links, we integrate multiomics data like TF-DNA-binding (Dibaeinia and Sinha 2020) in applications 6-7 to predict prior GRNs. We identify REs for TG and determine TFs likely to bind directly to or associate indirectly with TFBSs on these REs. We input *N* TG-specific candidate TFs to NetREm for TG. Then, we overlay NetREm’s *N*^*^ TF-TG regulatory links for TG with this prior GRN (initial TF–RE –TG links for *N* TFs for TG). This helps us annotate our links with epigenomic information on REs. Ultimately, we isolate highly-confident final TF-RE-TG links for our final *N*^*^ TFs.

#### Cell-type TF-TG regulatory network

We apply NetREm, iteratively, to each of the *G* TGs and weave together individual TF-TG links (details: **Supp. File 5**). We may narrow down links by retaining TFs with |*c*^*^| > *c*_*min*_ (min threshold, default: 0) and TGs meeting specific criteria (e.g. *MSE*_*TG*_ < *MSE*_*max*_). Our cell-type-specific complementary GRN relates TFs to TGs they regulate, helping explain how cell-types establish and maintain cellular identity. We may annotate/validate this network by identifying eSNPs impacting TF binding with eQTL links to altered TG expression (Coetzee et al. 2015); when prior GRNs are used, we ensure SNPs fall in the same REs where TFs are predicted to bind, linking them to TG regulation.

## Output 2: TF-TF coordination (direct/indirect interactions)

### TG-specific cell-type coordination *B*

NetREm helps fulfill the need for cell-type-specific proteome analysis by which proteins interact to carry out processes like TG regulation. Existing PPINs aggregate direct and/or indirect PPIs in an organism. This broad approach has limitations, as not all proteins are expressed in every cell or tissue type, and some may be aberrant in diseases (Padi and Quackenbush 2015). To overcome this, there are efforts to annotate global PPIs at various levels, including tissue-specific protein expression, cell-line-specific links, phenotype-based studies (e.g. CPPID) (Federico and Monti 2020). However, co-expression of TFs need not imply they interact in specific cell or tissue types (Gonzalez-Teran et al. 2022). Further, PPINs do not distinguish between cooperation and antagonism. This underscores the need for NetREm’s 2^nd^ output that predicts how TFs coordinate to regulate TG in the cell-type.

NetREm’s 2^nd^ output is a weighted and signed TG-specific TF-TF coordination network given by an adjacency matrix of coordination scores *B*, if *N*^*^ ≥ 2 final TFs for TG. These scores are a function of both embeddings 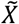 and *c*^*^: 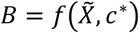. In our framework, *B*_*ij*_ > 0 suggests cooperativity (e.g. co-binding, pioneer-settler TF relations) and *B*_*ij*_ < 0 indicates antagonism (e.g. sequestration) between *TF*_*i*_ and *TF*_*j*_ for co-regulating TG.

We use *c*^*^ to predict the nature of interactions among *N*^*^ TFs for TG in a symmetric matrix *C* ∈ ℝ^*N* ×*N*^. Here, *C*_*ij*_ = *sign*(*c*_*i*_, *c*_*j*_) = {1 *i*f 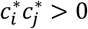; −1 if 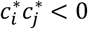; 0 otherw*i*se} and *C*_*ii*_ = 0. RTNduals assesses coordinated behavior of 2 TFs by analyzing correlation distributions between them and their shared TGs; building on approaches like these, we use *C* to deduce relative coordination relations among TFs for TG, acknowledging TFs may exhibit antagonistic or cooperative interactions depending on TG and context. If *C*_*ij*_ > 0, both TFs likely cooperate, aiming to either upregulate or downregulate TG expression in unison; their combined synergistic net effect on the TG is stronger than their individual effects. Conversely, if *C*_*ij*_ < 0, both likely act antagonistically, with conflicting influences on TG expression; this activator-repressor antagonism weakens their combined effect compared to their individual impacts, potentially due to partially canceling each other’s activities (Berenson et al. 2023). When *C*_*ij*_ = 0 (and *i* ≠ *j*), at least 1 of the 2 TFs is not a final TF and we cannot ascertain their potential nature of interaction.

Earlier, we set 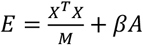. The 1^st^ term 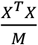 represents the original normalized inner product space from column vectors *x*_1_, *x*_2_, …, *x*_*N*_ of *X*, where *x*_*i*_ ∈ *R*^*M*^ represents *TF*_*i*_’s standardized expression levels across *M* cells in cell type. The 2^nd^ term *A* = *D*^*T*^(*W⨀V*)*D* purely depends on TF-TF PPIN strengths *w* > 0 and can be retrieved from public databases. By *Eq* (3*a*), 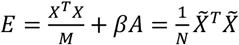; we thus transform *X* and network-prior PPIN data to 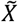 embedding data. This yields a new normalized inner-product space 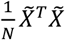 that helps depict and encode an aspect of cell-type TF-TF coordination scores for regulating TG. Since *N* is a scalar, we use 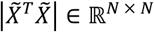. For each *TF*_*i*_ − *TF*_*j*_ pair, we divide 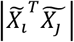, which is proportional to the extent of their potential coordination, by 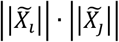 to scale it. 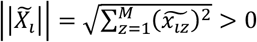 is the Euclidean norm of *TF*_*i*_’s embedding 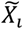. This essentially is their cosine similarity (cos) magnitude: 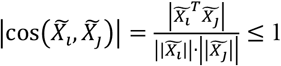. To learn coordination scores, we use coefficient-aware-cos metric: 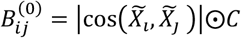 for *i* ≠ *j* and 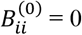. We apply max absolute value scaling 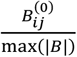 where max(|*B*|) is the max magnitude of 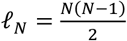 scores. Our TG-specific TF-TF coordination *B* ∈ ℝ^*N* × *N*^ has: 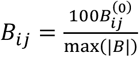 where −100 ≤ *B*_*ij*_ ≤ 100 if *TF*_*i*_ and *TF*_*j*_ are among *N* ^*^ TFs (i.e. 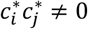), else 0. Of ℓ_*N*_ scores, 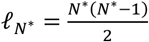 are ≠0.

TFs with higher |*B*| have stronger coordination for co-regulating TG. NetREm predicts *B* for known TF-TF PPIs (pairs with *w*_*ij*_ > η), uncovering meaningful, cell-type-specific PPI subnetworks. These documented PPIs have established partnership for orchestrating biological processes. It also predicts *B* for novel, artificial PPI links (*w*_*ij*_ = η), flagging (high |*B*|) promising FN TF-TF links for follow-up investigation. TF-TF coordination can be direct (e.g. form complexes, tethering) or indirect (e.g. TFs may not interact physically but can modify local chromatin environments, facilitating binding of other TFs) (Srivastava and Mahony 2020). Typically, direct PPIs rely on genome-wide data, indirect (e.g. guilt-by-association) PPIs use genetic interaction data (Wang et al. 2009). While studies predict negatomes (proteins unlikely to interact physically) (Jha et al. 2022), they offer limited insights on indirect PPIs.

### Cell-type coordination 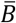

We define 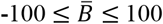 (math details in **Supp. File 5**). Each non-zero score in 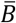 is unique. 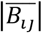 denotes relative strength of *TF*_*i*_ − *TF*_*j*_ coordination and cell-type co-regulation (i.e. normalized interaction significance); the sign encapsulates their net observed behavior across *G* TGs: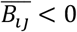: antagonistic, 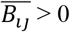: cooperate.

### Real-world Datasets & Pre-processing

We apply NetREm for each TG in the cell-type for 7 main applications spanning 2 organisms (humans: 1, 2, 5-7; mouse: 3, 4). We run NetREm without prior GRNs for these 5 applications: **1**. Simulated hESC data. **2**. HSCs: self-renewing, long-lived cells in bone marrow that help produce blood cells. **3**. mESCs: derived from inner cell mass of early embryo; they are pluripotent as they can self-renew, develop, specialize, differentiate, and mature into any cell type. **4**. mDCs: immune cells that capture and present antigens to other immune cells. **5**. PBMCs containing 9 sub cell-types. In these 5 applications, we fix *N* = 𝒩 candidate TFs for all TGs. If the TG is also a TF, we remove it from the set for the TG, so *N* = 𝒩 − 1. In applications 1-4, we use gold standards to hone our TGs and TFs so TF-TG regulatory links are comparable with ground truth (McCalla et al. 2023); we train models for TGs and for 𝒩 TFs found in both gene expression and ground truth data. For PBMCs, we train NetREm on 9 cell types with 1,029 TFs (Lambert et al. 2018). We run NetREm on 2 applications using prior GRNs from multi-omics to define a custom set of *N* highly probable, candidate TFs for each TG (*N* differs across TGs): **5**. mSCs vs. nmSCs in PNS. **6**. AD and Control stages for 8 CNS cell-types. 4 Glial cells: Astrocytes, Oligo, OPCs, Mic; 2 Neurons: GABA-ergic InNs and Glutamatergic Excitatory Neurons (ExNs); 2 Vascular and Blood-Brain Barrier (BBB) cells: Pericytes, Endothelial (Endo.) BBB cells. **Table S16-17** breaks down the # of TGs, # of TFs if *N* is fixed (applications 1 – 5) or metrics if variable (applications 6 – 7). For instance, in hESCs, we run NetREm 1,250 times (1 for each TG), with *N* = 206-207 TFs for each TG. **Supp. File 5** details parameters and evaluation for our applications.

### Single-cell gene expression data

**Supp. File 5** provides preprocessing details for our 7 main applications and additional datasets. **1:** We randomly select 1,250 TGs and corresponding TFs from weighted and signed (+: activates; -: represses) ground truth GRN atlas from TF induction analysis (Sharov et al. 2022). This results in 𝒩 = 207 TFs and 5,050 GRN links we input to SERGIO to simulate realistic single-cell data for 100 and 1k cells and 1,442 genes. We vary noise (30, 60, 90)%, retrieving 3 different synthetic expression datasets. **2:** We use (Buenrostro et al. 2018). **3:** We use (Tran et. al 2019) that reprograms mouse embryonic fibroblasts to embryonic-like induced pluripotent stem cells. **4:** We use normalized data (McCalla et al. 2023) from (Shalek et. al 2014) for >1.7k primary bone marrow DCs. **5:** We use public healthy donor data on 2.7k PBMCs based on (Satija et. al. 2024). **6:** We use (Avraham et al. 2022) for mSCs and nmSCs in DRG L4,5 regions from 5 donors. **7:** We use processed (Gupta et al. 2022) for 24 AD and 24 healthy humans for 8 cell types based on 80,660 droplet-based single-nucleus prefrontal cortex transcriptomes (Mathys et. al 2019).

### Prior GRN reference info on TG regulation

In applications 6-7, we employ multi-omics and epigenetic data from open scATAC-seq chromatin regions mapped to TGs (i.e. peak-TG links) to identify potential interacting REs for TGs. We map sequence-specific TF motifs to REs, using Position Weight Matrix (PWM) databases to predict TFBSs, forming a motif-based GRN: direct TF-RE-TG candidate links. We prune this motif-based TF list based on relative TF expression (proxy for protein abundance) and motif matching scores. To capture overlooked TFs, we augment our pruned TF list by adding TFs with known PPIs and predictions of TF-TF colocalization, complexes, and/or Molecular Function similarity (Wu et al. 2021). This addresses limitations of GRN-inference tools that rely solely on accessible motif matches and may miss causal TFs for TGs (Zhang et al. 2023); as ∼10% of the ∼1.6k human TFs lack motif data and are traditionally excluded, using PPIs during GRN inference is recommended to incorporate these missing TFs (Badia-i-Mompel et al. 2023). Augmented TFs may bind to TFBSs directly (weak signals) or indirectly (associate with DNA-binding TFs) (Gordân et al. 2009; Sloan et al. 2016). Our final prior GRNs comprise these initial TF-RE-TG links. We input the *N* biologically-promising TG-specific candidate TFs to NetREm for the given TG. These subjective steps are detailed in **Supp. File 5**.

### Organism-specific PPINs

We employ public STRINGdb (Szklarczyk et al. 2023) to construct human and mouse PPINs (details: **Supp. File 5**; we add more resources for humans) for proteins that partner through direct (physical binding, complex coexistence) and/or indirect (e.g., metabolic/signaling paths) PPIs (Oughtred et al. 2021). We scale average combined scores (many evidence types) to assign weights: 0.01 < *w* ≤ 1. Any self-loops are ignored. For each TG in a context, we filter this partly-connected PPIN 𝕎 to only keep its *N* TFs to yield *W*^(0)^. We run NetREm, assigning an artificial weight η = 0.01 to missing edges so 0.01 ≤ *w* ≤ 1 for *i* ≠ *j* to ensure numerical stability and propel discovery of novel TF-TF links; this yields our TG-specific final comprehensive fully-connected TF-TF PPIN (input *W*).

## Data access

We implement NetREm as an open-source software package: GitHub.com/SaniyaKhullar/NetREm with details in **Supp. File 5. Table S18** lists resources. We use human hg38 and rat rn5 reference genomes.

## Acknowledgements

D.W. conceived and designed the study. S.K., X.H., and D.W. developed the methodology for NetREm. In particular, X.H. primarily formulated the mathematical models in Steps 1 and 2, added the adaptive pseudo-inverses adjustment, and verified the linear algebra used in the method. X.H. and S.K. provided mathematical justification for Steps 1 and 2. S.K. and X.H. designed methodology for TF-TF coordination scores (from network regression embeddings) and TF-TG regulatory networks and designed simulation studies. S.K., X.H., R.R., J.S., and D.W. contributed to writing the manuscript. S.K. performed benchmarking, computational experiments, bioinformatics analysis, and created all supplementary materials. S.K. and X.H. wrote open-source code for NetREm. R.R. and J.S. conducted experiments in rats, providing external data and analysis for Schwann cell validation. S.K., J.S., and D.W. analyzed the data. All authors have read and approved the final manuscript and are grateful to receive funding from these sources: National Institutes of Health, R01AG067025, RF1MH128695, R03NS123969, R21NS127432, R21NS128761 to D.W. and P50HD105353 to Waisman Center. S.K. is supported by an NLM training grant to Computation and Informatics in Biology and Medicine Training Program (NLM 5T15LM007359). Authors are grateful for feedback and advice from Kalpana Hanthanan Arachchilage, Ph.D., Jerome Choi, Noah Cohen Kalafut, Sean Chang, and Harrison Pantera. In addition, the authors appreciate feedback from Renu Poochie Khullar and Armaan Khullar for the manuscript.

